# Dysfunction of exocytosis causes catecholamine hypersecretion in patient with pheochromocytoma

**DOI:** 10.1101/2021.11.02.466874

**Authors:** Sébastien Houy, Laura Streit, Ines Drissa, Marion Rame, Charles Decraene, Sophie Moog, Laurent Brunaud, Joël Lanoix, Rabie Chelbi, Florence Bihain, Stéphanie Lacomme, Sandra Lomazzi, Michel Vix, Didier Mutter, Eustache Paramithiotis, Christophe Dubessy, Nicolas Vitale, Stéphane Ory, Stéphane Gasman

## Abstract

Pheochromocytoma (Pheo) is a neuroendocrine tumor that develops from chromaffin cells of the adrenal medulla, and is responsible of an excess of catecholamines secretion leading to severe clinical symptoms such as hypertension, elevated stroke risk and various cardiovascular complications. Surprisingly, hypersecretory activity of Pheo has never been explored at the cellular and molecular levels from individual tumor cells. In the present study, we have combined catecholamine secretion measurement by carbon fiber amperometry on human tumor cells directly cultured from freshly resected Pheo, with the analysis by mass spectrometry of the exocytotic proteins differentially expressed between the tumor and the matched adjacent non-tumor tissue. Catecholamine secretion recordings from individual Pheo cells obtained from most patients reveal a higher number of exocytic events per cell associated with faster kinetic parameters. Accordingly, we unravel significant tumor-associated modifications in the expression of key proteins involved in different steps of the calcium-regulated exocytic pathway. Altogether, our findings indicate that dysfunction of the calcium-regulated exocytosis at the level of individual Pheo cell is a cause of the tumor-associated hypersecretion of catecholamines.

## Introduction

Through the secretion of hormones and neuropeptides, the neuroendocrine system controls many vital functions such as metabolism, blood pressure, reproduction, growth and development, stress and eating behavior, to cite only a few. Neoplasms can derive from all kinds of hormone secreting cells giving rise to a neuroendocrine tumor (NET). NETs constitute a highly heterogeneous group of neoplasm, but share a common feature in that they are often associated with a deregulation of hormone secretion, mainly hypersecretion, which can induce symptoms and major clinical complications (Zandee *et al*, 2017). Therefore, the secretory pathways and their dysfunction appear as an important issue to be considered in NETs. However, the cellular and molecular mechanisms underlying hypersecretory activity of NETs remain poorly known.

In neuroendocrine cells, hormones and neuropeptides are stored in large dense core vesicles (secretory granules) and are secreted through calcium-regulated exocytosis, a process that has been intensively studied for decades (Anantharam & Kreutzberger, 2019). It involves several tightly regulated trafficking steps including the recruitment of secretory granules to the cell periphery, their docking to exocytic sites of the plasma membrane, their priming and finally the fusion between the secretory granule membrane and the plasma membrane leading to the release of the intra-granular content (Burgoyne & Morgan, 2003; Lang & Jahn, 2008). Chromaffin cells of the adrenal medulla, which store and then secrete catecholamines into the blood stream, have been widely used by us and others as an experimental model to uncover the molecular mechanisms controlling calcium-regulated exocytosis (Bader *et al*, 2002; Gasman & Vitale, 2017; Malacombe *et al*, 2006).

The NETs deriving from chromaffin cells of the adrenal medulla are called pheochromocytomas (Pheo) (Lenders *et al*, 2020). Most of the Pheos are responsible for catecholamine hypersecretion leading to clinical symptoms such as permanent or paroxysmal hypertension or to cardiovascular complications including myocarditis, Takotsubo syndrome and various forms of cardiomyopathies (Lenders *et al*., 2020; Pappachan *et al*, 2018; Pourian *et al*, 2015; Zhang *et al*, 2017). The reason for this excess of secretion is currently not known. Among the likely possibilities are an anarchic proliferation of secretory cells or an intensification of the secretory capacity at the single cell level. We therefore attempted to investigate the cellular mechanisms responsible for a possible specific secretion dysfunction in Pheo by performing carbon fiber amperometry on primary culture of human Pheo cells. This technique allows the precise measurement of individual exocytotic event dynamics in real time and in single cells (Fathali & Cans, 2018; Mosharov & Sulzer, 2005). Combined with the detection in human Pheo tissue of exocytotic protein expression changes by mass spectrometry, it allowed us to reveal upregulated exocytosis at the single cell level and to identify specific steps of the exocytotic process that are dysregulated in the tumor as well as potential actors of the molecular machinery triggering hypersecretion in Pheo.

## Results

### Technical workflow and overall patient characteristics

This study includes two types of analyses (see technical workflow in Figure 1) performed on histologically confirmed Pheo samples from 27 distinct patients. On one side, we have analyzed the secretory activity of single tumor cells. To do so, freshly resected Pheos originating from 22 patients were placed in primary culture in order to perform real-time single cell catecholamine secretion measurement using carbon fiber amperometry (Figure 1A). On the other side, we have used quantitative mass spectrometry analysis to measure the relative differential expression of proteins involved in the exocytic pathway from 5 other Pheo tissues, which were compared to the matched patient non-tumor adrenal tissue. To conduct this proteomic analysis, two subcellular fractions enriched for exocytic proteins (the cytosolic fraction and a low-density membrane fraction containing plasma membrane, Golgi, endosomes and secretory vesicles) were isolated from the pairs of matched non-tumor and tumor frozen tissues (Figure 1B). Note that due to the low amount of human material, it is technically impossible to perform both amperometry and mass spectrometry experiments on the same samples.

**Figure 1:**
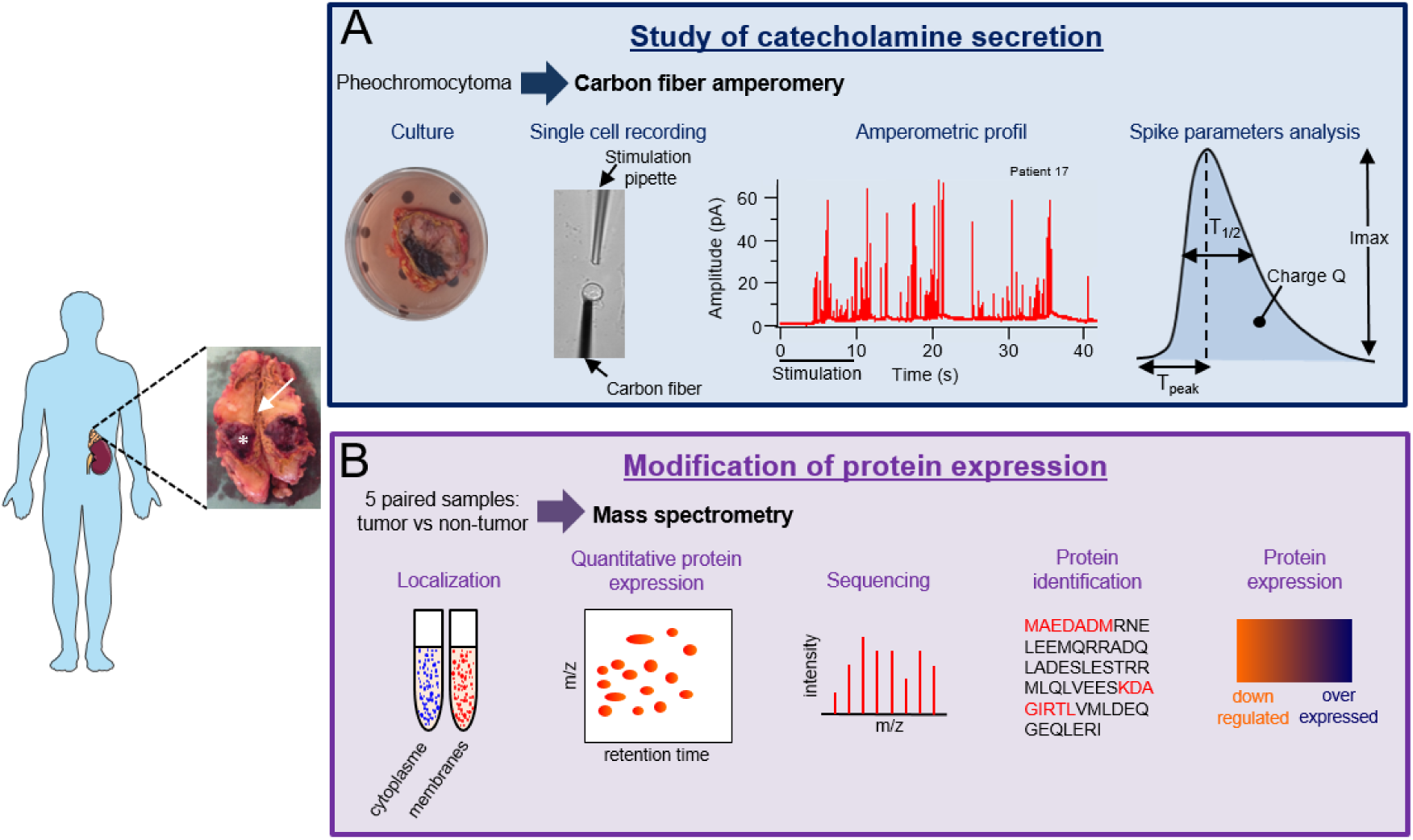
Technical workflow of catecholamine secretion measurement and comparative proteomic analysis of human Pheo. A resected adrenal gland, cut in half, from a patient with Pheo is shown (asterisk). The non-tumor tissue is shown by the arrow. (A) Description of the different steps of the catecholamine secretion measurement by carbon fiber amperometry from the primary cell culture of the tumor to the amperometric spike analysis. A representative amperometric trace of a Pheo cell is illustrated. The dark bar indicates when a 100 µM nicotine solution was applied. The parameters of individual spike that were analyzed are indicated. (B) Complete protein profiling workflow of differential mass spectrometry analysis between Pheo tissue and matched non-tumor tissue.

For patients included in the amperometric analysis, a slight predominance of male was found (59%) with a mean age of 50.5 ± 10 years at diagnosis (Table 1). Sixteen patients (73%) were diagnosed with hormonal-related symptoms whereas all of them presented abnormal hormonal secretion (100%) including adrenergic or noradrenergic phenotype in 13 (59%) and 9 (41%) cases, respectively. Seven patients (46%) out of 16 tested were diagnosed with a genetic predisposition (4 *NF1*, 1 *RET*, 1 *SDHB* and 1 *SDHD*). The mean tumor size was 4.5 cm (range 1.7-8 cm). Other biological and clinical characteristics are detailed in Table 1.

**Table 1:** Biological and clinical characteristics of the 22 patients from which Pheo were used for amperometric analysis.

| Characteristics, *n* available |  |
| --- | --- |
| **Age at diagnosis** (mean ± SD, years), *n*=22 | **50.5 ± 10** |
| **Males** (%), *n*=22 | **13 (59%)** |
| **Symptoms at diagnosis:** *n*=22 |  |
| - Tumor-related symptoms (%) | **4 (18%)** |
| - Hormonal-related symptoms (%) | **16 (73%)** |
| **Hormonal hypersecretion** |  |
| - Adrenergic/noradrenergic phenotype, *n*=22 | **13/9** |
| - Urinary MN, ULN ratio (mean ± SD), *n*=19 | **9.3 ± 14** |
| - Urinary NMN, ULN ratio (mean ± SD), *n*=19 | **7.7 ± 5.6** |
| - Plasma free MN, ULN ratio (mean ± SD), *n*=15 | **4.5 ± 5.0** |
| - Plasma free NMN, ULN ratio (mean ± SD), *n*=15 | **6.2 ± 6.1** |
| - Plasma CGA, ULN ratio (mean ± SD), *n*=12 | **3.4 ± 2.6** |
| **Pathology:** |  |
| - Tumor size (mean + range), *n*=21 | **4.5 (1.7-8)** |
| - Ki-67 (mean + range), *n*=12 | **2.1 (1-5)** |
| - PASS score (mean + range), *n*=22 | **1.9 (0-6)** |
| **Genetics:** *n*=16 |  |
| - No germline mutation (%) | **9 (56.25%)** |
| - *NF1* | **4 (25%)** |
| - *RET* | **1 (6.25%)** |
| - *SDHB* | **1 (6.25%)** |
| - *SDHD* | **1 (6.25%)** |

*MN: metanephrine, NMN: normetanephrine, ULN: upper limit normal, CGA: chromogranin A, PASS: Pheochromocytoma of the Adrenal Gland Scaled Score, NF1: Neurofibromatosis type 1, RET: Rearranged during transfection, SDHB: Succinate dehydrogenase B, SDHD: Succinate dehydrogenase D.*

Biological and clinical characteristics of the 5 patients included for the proteomic analysis are detailed in Table 2. With a mean age of 58 ± 8 years, these 5 patients had hormonal-related symptoms at diagnosis, four of which were classified as adrenergic phenotype and one as noradrenergic phenotype. None of the patients had germline mutation (out of 3 patients tested). The mean tumor size was 5.3 cm (range 3-8 cm). Individual characteristics of the 27 patients can be found in Supplementary Table 1.

**Table 2:** Biological and clinical characteristics of the 5 patients from which Pheo were used for proteomic analysis.

| Characteristics, *n* available |  |
| --- | --- |
| **Age at diagnosis** (mean ± SD, years), *n*=5 | **58 ± 8** |
| **Males** (%), *n*=5 | **2 (40%)** |
| **Symptoms at diagnosis:** *n*=5 |  |
| - Tumoral-related symptoms (%) | **1 (20%)** |
| - Hormonal-related symptoms (%) | **5 (100%)** |
| **Hormonal hypersecretion** |  |
| - Adrenergic/noradrenergic phenotype *n*=5 | **4/1** |
| - Urinary MN, ULN ratio (mean ± SD), *n*=4 | **3.7 ± 4.2** |
| - Urinary NMN, ULN ratio (mean ± SD), *n*=4 | **6.8 ± 8** |
| - Plasma free MN, ULN ratio (mean ± SD), *n*=2 | **3.0 ± 2.8** |
| - Plasma free NMN, ULN ratio (mean ± SD), *n*=2 | **3.0 ± 2.8** |
| - Plasma CGA, ULN ratio (mean ± SD), *n*=2 | **4.7 ± 2.9** |
| **Pathology:** |  |
| - Tumor size, cm (mean + range), *n*=5 | **5.3 (3-8)** |
| - Ki-67, % (mean + range), *n*=3 | **3.3 (1-7)** |
| - PASS score (mean + range), *n*=3 | **3 (0-9)** |
| **Genetics:** *n*=3 |  |
| - No germline mutation (%) | **3 (100%)** |

*MN: metanephrine, NMN: normetanephrine, ULN: upper limit normal, CGA: chromogranin A, PASS: Pheochromocytoma of the Adrenal Gland Scaled Score.*

### Analysis of catecholamine secretion in human pheochromocytoma by carbon fiber amperometry

A representative amperometric trace recorded from a human Pheo cell is illustrated in Figure 1A. Each individual spike represents a single granule fusion event and is composed of a rapid rise of the electrode current corresponding to the oxidation of catecholamines quickly released at high concentration through the fusion pore as it dilates. The spike rise is then followed by a slower decay representing a decreased of the catecholamine flux through the pore as the granule empties. In addition to the quantification of number of events per cell, the analysis of individual amperometric spikes provides valuable dynamic information on the exocytic process. Hence, the surface area or quantal size (Q) is proportional to the amount of catecholamines released per event, the spike amplitude value (Imax) reflects the maximal flux of catecholamines, whereas the half-width (*T*_1/2_), and the time to peak (*T*_*peak*_) reflect the duration of the exocytotic event and the kinetics of the fusion pore expansion, respectively (Figure 1A).

Primary culture of human Pheo cells is rather efficient as most attempts were successful. However, culturing non-tumor chromaffin cells taken outside the tumor zone sample appeared trickier and failed most of the time for unidentified reasons. Nevertheless, we were able to obtain 4 different cultures of non-tumor cells that could be used for amperometric recordings. Therefore, we have compared the distribution of the amperometric parameters of each of the 22 patients individually with the mean values calculated from these 4 non-tumor samples. All the amperometric parameters are detailed in Table 3. The major change concerns the total number of spikes. Indeed, among the 22 patients, 16 (73 %) exhibit a significant increase of the number of spikes (11 patients with an increase up to 2 fold and 5 patients with an increase above 2 and up to 3.4 fold; Figure 2). Hence these data suggest that one of the main causes of tumor-associated catecholamine hypersecretion could be an increase of the number of exocytic events.

**Table 3:** Characteristics of amperometric spikes from 22 Pheos and 4 non-tumor tissues.

| Identification | Number of cells | Total number of spikes | Total number of spikes/cell | Number of spikes | I<sub>max</sub> (pA) | Charge (pC) | T<sub>1/2</sub> (ms) | T<sub>peak</sub> (ms) |
| --- | --- | --- | --- | --- | --- | --- | --- | --- |
| Non-tumor 1 | 19 | 224 | 11.79 ± 0.67 | 146 | 41.56 ± 2.23 | 2.22 ± 0.13 | 51.90 ± 1.14 | 42.21 ± 1.24 |
| Non-tumor 2 | 11 | 164 | 14.91 ± 2.05 | 126 | 10.92 ± 0.81 | 0.92 ± 0.10 | 68.41 ± 3.41 | 29.87 ± 1.67 |
| Non-tumor 3 | 8 | 144 | 18.00 ± 3.44 | 102 | 18.77 ± 2.35 | 0.73 ± 0.20 | 30.76 ± 3.94 | 13.50 ± 2.07 |
| Non-tumor 4 | 6 | 214 | 35.67 ± 2.17 | 112 | 15.51 ± 1.65 | 0.97 ± 0.12 | 56.13 ± 4.26 | 28.50 ± 2.10 |
| Patient 1 | 19 | 609 | **32.05 ± 6.61 \*\*** | 469 | 21.34 ± 2.16 | **0.90 ± 0.10 \*** | **38.97 ± 2.70 \*\*\*** | **16.37 ± 1.41 \*\*\*** |
| Patient 2 | 22 | 653 | **29.68 ± 3.26 \*\*\*** | 518 | **12.45 ± 0.65 \*\*\*** | **0.71 ± 0.04 \*\*\*** | 52.11 ± 1.69 | **23.69 ± 1.03 \*\*\*** |
| Patient 3 | 24 | 1388 | **57.83 ± 8.01 \*\*\*** | 1032 | 27.85 ± 2.91 | 1.79 ± 0.16 | **61.69 ± 2.74 \*** | 29.26 ± 1.42 |
| Patient 4 | 4 | 39 | 9.75 ± 1.70 | 38 | 13.08 ± 3.20 | 0.71 ± 0.26 | 44.90 ± 8.53 | 21.05 ± 4.65 |
| Patient 5 | 12 | 326 | **27.17 ± 4.74 \*** | 240 | **13.82 ± 0.92 \*** | 1.30 ± 0.11 | **80.38 ± 5.65 \*\*\*** | 36.49 ± 2.11 |
| Patient 6 | 18 | 499 | **27.72 ± 1.60 \*\*\*** | 320 | **45.78 ± 2.00 \*\*\*** | **2.33 ± 0.10 \*\*\*** | **48.34 ± 0.76 \*** | 40.43 ± 0.90 |
| Patient 7 | 17 | 524 | **30.82 ± 3.32 \*\*\*** | 408 | **11.04 ± 0.58 \*\*** | 0.96 ± 0.07 | **69.70 ± 3.27 \*\*** | 30.43 ± 1.43 |
| Patient 8 | 6 | 185 | **30.83 ± 5.46 \*\*** | 163 | 30.99 ± 4.50 | 2.00 ± 0.29 | 65.30 ± 6.54 | 31.31 ± 2.67 |
| Patient 9 | 31 | 682 | **22.00 ± 1.85 \*** | 416 | **40.28 ± 1.61 \*\*\*** | 1.71 ± 0.08 | **40.29 ± 1.13 \*\*\*** | 34.45 ± 1.18 |
| Patient 10 | 59 | 1382 | **23.42 ± 2.04 <sup>T</sup>** | 1231 | 21.26 ± 1.28 | 1.25 ± 0.08 | 52.81 ± 2.19 | **26.49 ± 1.06 \*\*** |
| Patient 11 | 12 | 161 | 13.42 ± 1.31 | 130 | 17.39 ± 2.35 | **0.85 ± 0.11 \*** | **42.54 ± 2.76 \*** | **19.00 ± 1.75 \*\*\*** |
| Patient 12 | 21 | 345 | 16.43 ± 2.01 | 292 | 20.17 ± 2.71 | **0.81 ± 0.10 \*\*** | **35.45 ± 1.20 \*\*\*** | **16.75 ± 0.84 \*\*\*** |
| Patient 13 | 23 | 428 | 18.61 ± 2.16 | 353 | **11.28 ± 0.81 \*\*\*** | **0.68 ± 0.05 \*\*\*** | 52.01 ± 2.47 | **25.47 ± 1.13 \*\*** |
| Patient 14 | 4 | 74 | 18.50 ± 5.24 | 59 | 27.09 ± 5.68 | 1.01 ± 0.14 | **35.08 ± 1.80 \*** | **16.11 ± 0.79 \*** |
| Patient 15 | 9 | 315 | **35.00 ± 6.88 \*\*** | 187 | **13.91 ± 1.93 \*** | **0.49 ± 0.08 \*\*\*** | **30.03 ± 1.99 \*\*\*** | **14.59 ± 1.13 \*\*\*** |
| Patient 16 | 12 | 482 | **37.08 ± 5.96 \*\*\*** | 334 | 21.58 ± 2.30 | **0.93 ± 0.14 \*** | **37.39 ± 2.83 \*\*** | **20.30 ± 3.34 \*\*** |
| Patient 17 | 15 | 866 | **57.73 ± 2.76 \*\*\*** | 400 | 23.72 ± 1.15 | 1.04 ± 0.06 | **39.65 ± 0.89 \*\*\*** | 30.66 ± 0.84 |
| Patient 18 | 9 | 266 | **29.56 ± 5.66 \*** | 140 | 19.18 ± 1.75 | 1.72 ± 0.24 | **79.57 ± 6.60 \*\*\*** | **41.97 ± 4.35 \*** |
| Patient 19 | 9 | 270 | **30.00 ± 5.31 \*\*** | 197 | **13.96 ± 1.03 \*** | **0.55 ± 0.04 \*\*\*** | **35.21 ± 1.19 \*\*\*** | **17.32 ± 0.74 \*\*\*** |
| Patient 20 | 9 | 379 | **42.11 ± 5.25 \*\*\*** | 276 | 15.84 ± 2.19 | **0.60 ± 0.10 \*\*\*** | **33.53 ± 2.29 \*\*\*** | **16.95 ± 1.58 \*\*\*** |
| Patient 21 | 4 | 73 | 18.25 ± 4.23 | 59 | 16.29 ± 5.83 | 0.74 ± 0.19 | 45.37 ± 2.51 | 22.65 ± 1.94 |
| Patient 22 | 5 | 151 | **30.20 ± 5.42 \*** | 82 | 25.13 ± 5.40 | 1.72 ± 0.30 | 65.82 ± 6.82 | 32.50 ± 3.74 |

*Amperometric recordings were performed on primary culture of human Pheo cells or non-tumor cells. Cells were stimulated with 100 μM of nicotine for 10 s. The number of cells, the number of amperometric spikes per cell and the different amperometric spikes parameters are indicated. Results represent the mean ± SEM. Bold values are considered significantly different from the control conditions; <sup>T</sup>p=0.05, \*p< 0.05, \*\*p< 0.01; \*\*\*p< 0.001 compared to the mean of non-tumor tissue; non-paired Wilcoxon test.*

**Figure 2:**
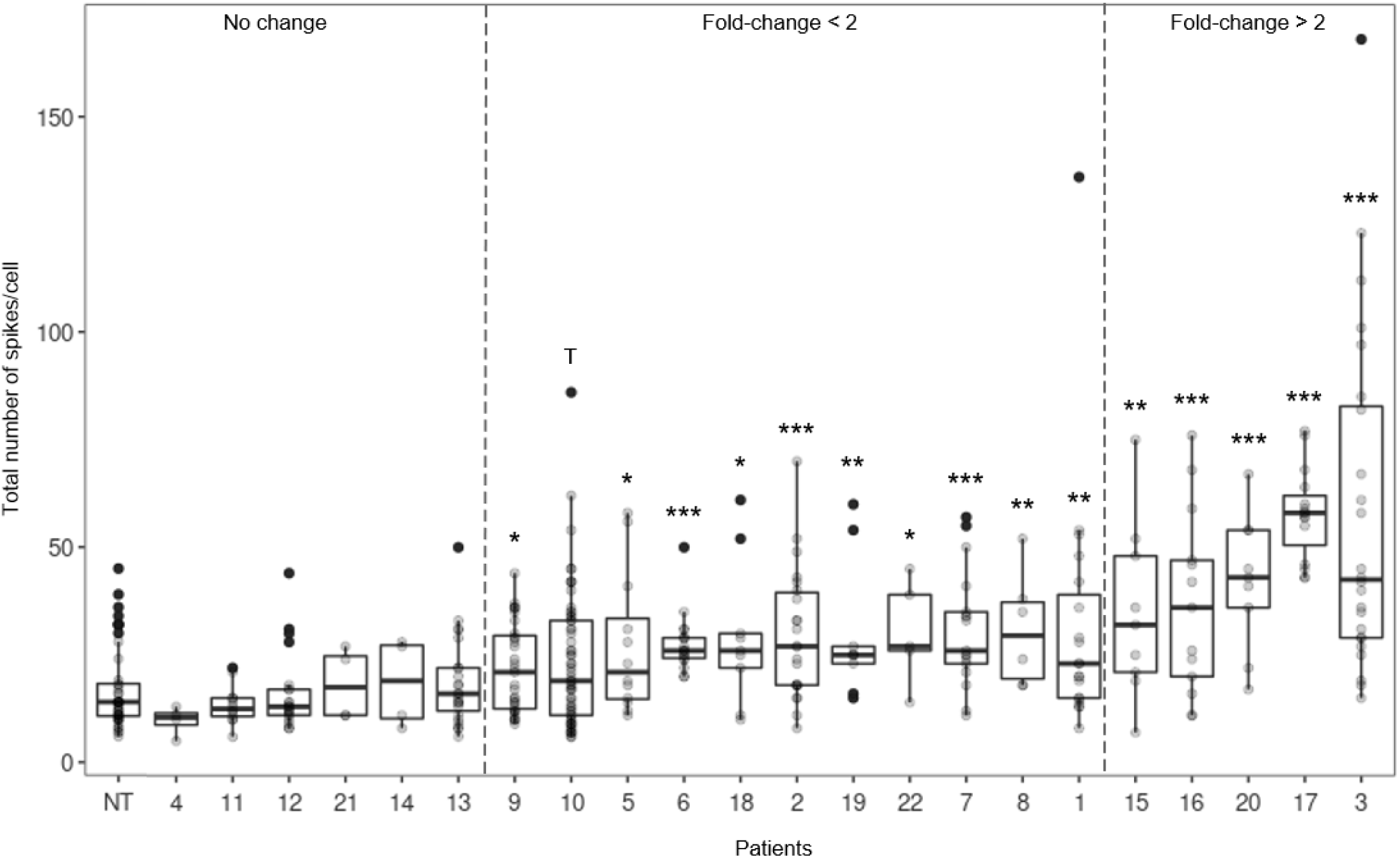
Analysis of catecholamine secretion events in Pheo cells from each patient by carbon fiber amperometry. Box- and-whisker diagrams illustrating the distribution of the number of amperometric spikes per cell for non-tumor cells (NT) and for cells from each patient with Pheo. Patients are classified according to the increasing effect of the number of spikes compared to non-tumor cells: no significant change, fold-change < 2 and fold-change > 2; ^T^p=0.05, *p< 0.05, **p< 0.01; ***p< 0.001 compared to the mean of non-tumor tissue; non-paired Wilcoxon test.

To better understand which parameters are mostly driving the secretory tumor profiles and to sort out potential patient clusters, we performed a principal component analysis (PCA) using the amperometric data obtained from the 22 tumor cell cultures (Figure 3). As shown in Figure 3A, the 3 first principal components explain all the data variations (97.3%). The correlation matrix indicated that the main amperometric parameters explaining the variation of the data within the 2 first dimensions are the charge and the *T*_*peak*_ and to a lesser extend the *T*_1/2_, parameters that are all correlated as shown by the variable projection analysis. The number of spikes per cell are less correlated and largely contributes to the third PCA dimension. Thus, it is interesting to note that the distribution of the patient samples plotted according to the first and the third component reveals a clear separation between populations with and without exocytic event increase as compared to non-tumor cells (Figure 3B). The distribution of the patient samples plotted according to the two first components forms one major cluster including most of the patients for which the two kinetic parameters, *T*_1/2_ and *T*_*peak*_, are significantly decreased (Figure 3C). Indeed, 14 (64%) patient’s tumor cell cultures presented a significant reduction of the *T*_1/2_ and/or *T*_*peak*_, often accompanied (9 patients) with a reduction of the quantal size (Charge Q; Figure 4A, B and Table 3). This type of amperometric profile corresponds to exocytic events occurring with a faster kinetics compared to normal cells. Interestingly, among these 14 patient samples, 10 displayed a concomitant increase of spikes per cells (Figure 2). Only 4 patient’s cultured cells (18%) displayed, on contrary, a significant increase of their *T*_1/2_ leading, for two patients, to a reduce spike amplitude (Imax, patients 5 and 7, Figure 4C and Table 3) and accordingly to a slower release kinetics. Finally, 4 patient’s cultured cells (18%) do not show significant changes in charge, *T*_1/2_ or T_*peak*_ indicating that the kinetic of the secretion in unaffected (Figure 4D). Altogether, our analysis of the amperometric recordings indicate that catecholamine secretion in tumor cells from patients with Pheo often involves a high number and fast secretory events, which most likely contribute to the tumor-associated hypersecretion.

**Figure 3:**
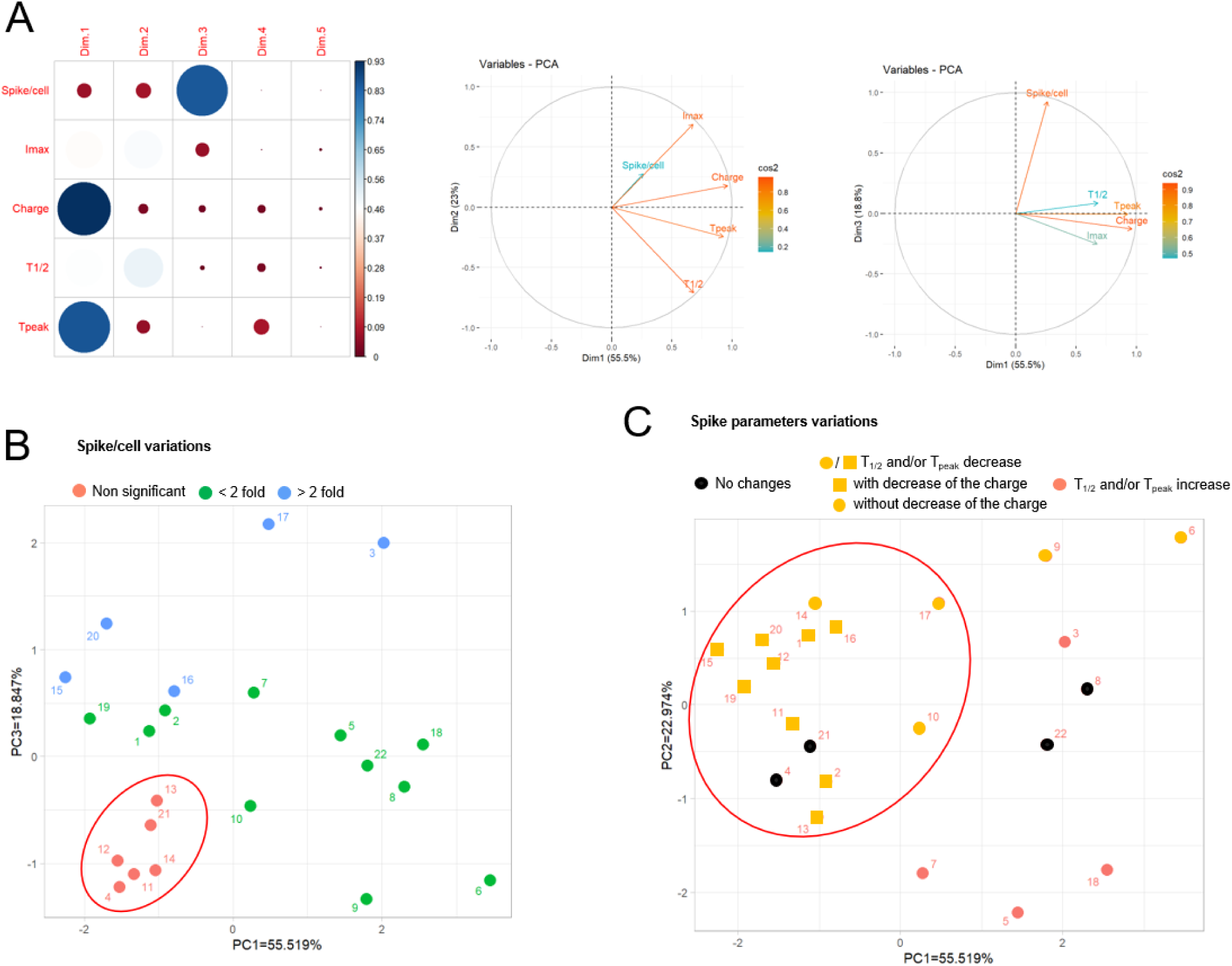
Principal component analysis (PCA) of amperometric spike parameters corresponding to catecholamine secretion recording on tumor cells cultured from 22 patients with Pheo. Number of spikes/cell and spike parameters (Imax, Charge, T_1/2_ and T_peak_) were subjected to PCA. (A) Correlation matrix and variable factor maps along the PCA dimensions. The first, second and third dimension (Dim) of principal components explain 55.5%, 23.0% and 18.8% of the data variations, respectively. The spike parameters Charge and T_peak_ largely contribute to Dim 1 whereas T_1/2_ and number of spikes per cell contribute to Dim 2 and 3, respectively. PCA variable vector map showing the projection of Imax, Charge, T_peak_ and T_1/2_ on Dim2 (right) and Dim3 (left). The projection of each variable vector gives an indication of the relation of these variables to T_peak_ and Charge (Dim1) or spikes/cell (Dim3) (B-C) The two-dimensional representation of PC1 and PC3 (B) allowed to separate patients according to the significant variation of the number of spikes recorded per cell (red circle = no changes; green circles = fold change < 2; blue circles = fold change > 2). The two-dimensional representation of PC1 and PC2 (C) revealed a large cluster of patients for which T_1/2_ and/or T_peak_ decreased (yellow squares and circles).

**Figure 4:**
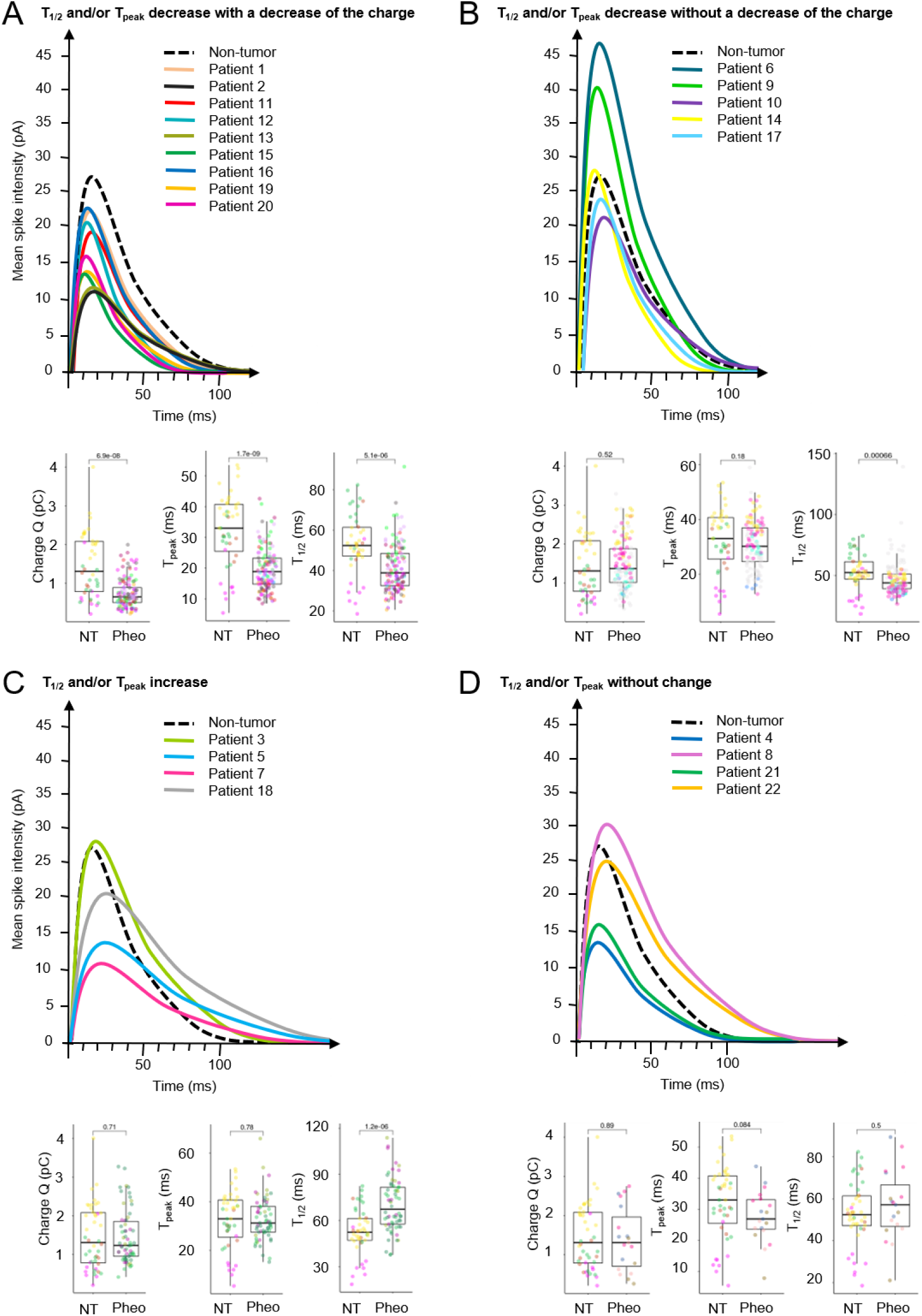
Analysis of amperometric spike parameters in Pheo cells from each patient. Superimposition of the averaged spike obtained for cells of each patient and distribution of the values corresponding to the spike charge (Q), the spike rise time (T_peak_) and the spike half-width T_1/2_ are shown for the 4 different populations of patients: decrease of T_1/2_ and/or T_peak_ with lower charge (A), decrease of T_1/2_ and/or T_peak_ without lower charge (B), increase of T_1/2_ and/or T_peak_ (C) and no significant changes of T_1/2_ and/or T_peak_ (D). Box- and-whisker diagrams illustrate the distribution of each variable values pooled from all the patients belonging to the indicated secretory profile. P values are indicated and patients are color coded. All the average amperometric data are detailed in Table 3.

### Differential protein expression of the exocytic machinery in human pheochromocytoma

Since catecholamine release is altered in Pheo, we asked whether proteins of the core machinery of secretory granule exocytic process could be deregulated. To do so, we performed a mass spectrometry analysis comparing 5 pairs of human Pheo with their respective matched adjacent non-tumor tissue. To increase detection and sensitivity, we conducted the proteomic analysis on purified subcellular fractions in place of total homogenate. Two main subcellular fractions were isolated from each tissue sample, a membrane fraction enriched for organelles and vesicles derived from the exocytic pathway and a fraction enriched in cytosolic proteins. From our proteomic data set, we selected proteins from the regulated exocytosis (#BPGO:0045055) and secretory granule (#CCGO:0030141) GO terms and focused on proteins significantly up or downregulated in the tumor by comparison with the corresponding non-tumor tissue, with fold change values greater than or equal to 2. In these conditions, we identified 166 deregulated proteins: 62 proteins from the cytosolic fraction, 54 from the membrane-enriched fraction and 50 common to both fractions (Figure 5A). Supplementary Table 2 details the median value of the relative expression changes of the 5 pairs of Pheo samples for all these 166 selected proteins associated with their known function. The volcano plot in Figure 5B shows the relationship between the p-values and the fold change in expression. The unsupervised hierarchical clustering of all differentially expressed proteins in row according to the paired samples in columns, is presented in Figure 5C for the corresponding cellular compartment. Note that the expression of proteins found in common in the membranous and the cytosolic fractions varies in the same direction (Supplementary Figure 1). This specific dataset of proteins corresponding to the proteins involved in the machinery of secretory granule exocytotic process clearly differentiate, by their modulation of expressions, the Pheo samples from the non-tumor samples. The functions of the deregulated proteins linked to secretory granule exocytosis include mainly: secretory granule organization and biogenesis, hormone processing, vesicular trafficking, docking, priming, membrane fusion, actin cytoskeleton organization, and small GTPases (Supplementary Table 2 and Figure 5D). Hence, we observed that numerous secretory granules cargos, as well as different proteins involved in the control of the frequency and the dynamic of the exocytic events are up-regulated.

**Figure 5:**
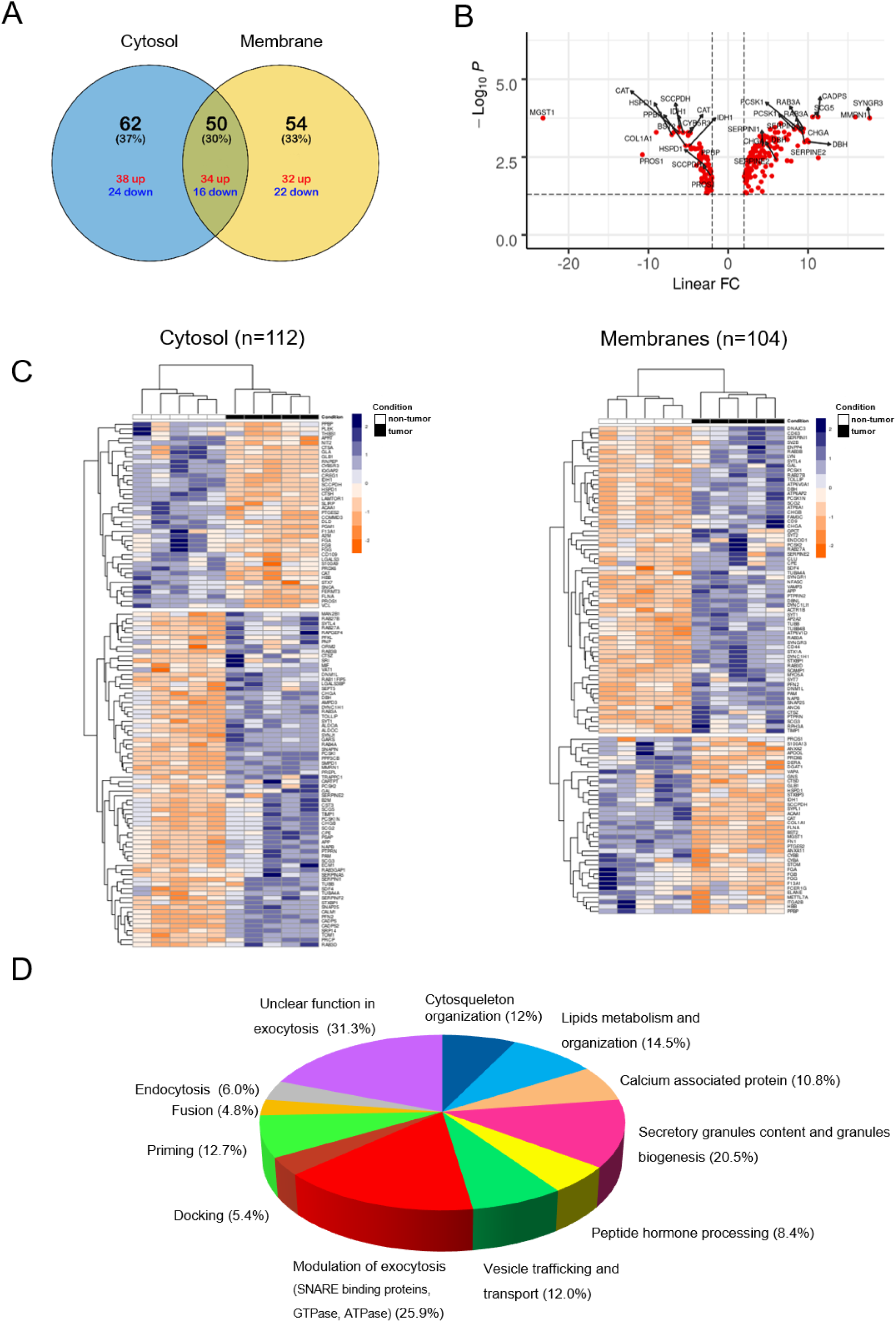
Identification of differentially expressed proteins involved in the exocytotic pathway in human Pheo tissue. (A) Venn diagram showing the distribution of the differentially expressed proteins in fractions enriched in cytosolic and membrane proteins. Colors indicate up-(red) or down-(blue) regulation. (B) Volcano plot of all differentially expressed proteins between Pheo and non-tumor samples. (C) Two-way hierarchical clustering of paired samples from each patient (non-tumor in white and tumor in black) in column according to the validated cytosolic (left) and membranous (right) proteins of the data set in row. Protein expression values were z-score normalized prior to clustering using the complete-linkage method together with the Euclidean distance. Each row represents a differentially expressed protein and each column, a patient according to the tissue type. The color scale illustrates the relative level of protein expression: red, higher expression; blue, lower expression. (D) Functional enrichment analysis of identified proteins. Each differentially expressed protein selected for their potential role in calcium-regulated exocytosis (Table 4) has been classified in one or several indicated biological functions.

To further assess the relevance of our data, we have compared our proteome dataset to a published gene expression analysis of 60 Pheo and 6 normal adrenal medulla tissues ((Lopez-Jimenez *et al*, 2010) https://www.ncbi.nlm.nih.gov/geo/query/acc.cgi?acc=GSE19422). Using this analysis, we have extracted 170 genes that belong at least to one of the 2 GO terms used in the proteomic analysis (regulated exocytosis and secretory granules) and that are significantly differentially expressed by at least two-fold between Pheo and normal adrenal tissues. Interestingly, among these 170 genes, 59 are common to our proteome dataset. As observed at the level of the protein expression changes, the hierarchical clustering of these 59 differentially expressed genes efficiently separated the tumor samples from the normal adrenal tissue (Figure 6A). Moreover, the expression of 53 of these 59 genes (90%) shows the same variation trend as the protein expression, both in the cytosol and in the membranes (Figures 6B and Supplementary Figure 2). To visualize whether the deregulated proteins identified in this comparison were highly connected to each other, we performed a protein-protein interaction (PPI) network analysis using Cytoscape ((Shannon *et al*, 2003), Figure 7). Interestingly, one major significant cluster is composed of proteins that are overexpressed both at the protein and at the gene levels (blue squares). The cluster includes proteins involved in the secretory granule composition such as various chromogranins (CHGA, CHGB, SCG2, SCG3, SCG5), in catecholamine synthesis or in hormone processing (DBH, PCSK1, PCSK2, CPE, PAM), in granule docking and fusion such as different SNAREs or SNAREs interacting proteins (SNAP25, STX1A, SYTL4, STXBP1, SYT1) and Rab GTPases (RAB3A, RAB3D, RAB27B).

**Figure 6:**
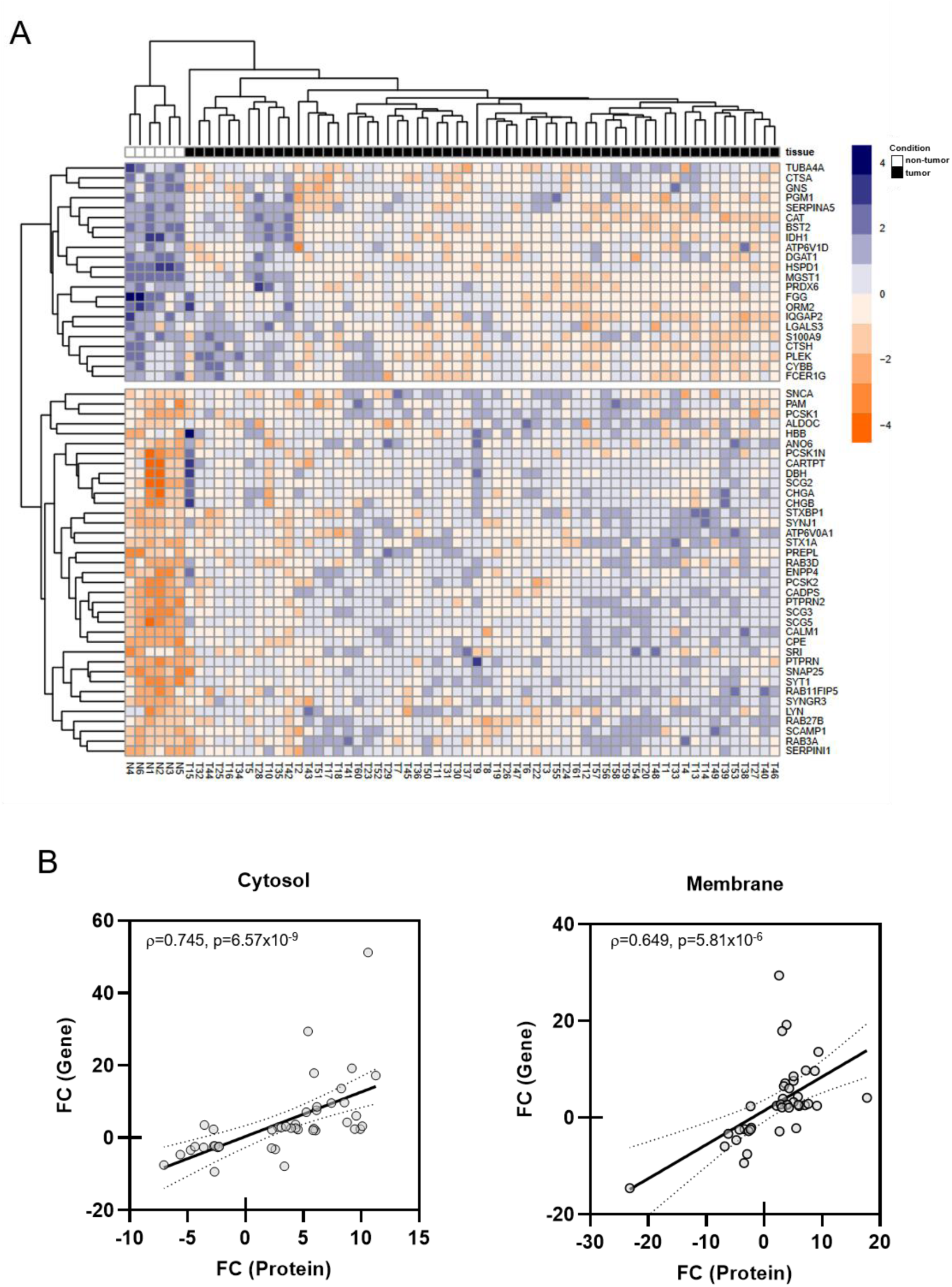
Comparison of the proteome dataset with a published gene expression analysis of 60 Pheo and 6 normal adrenal medulla tissues. (A) Unsupervised hierarchical clustering of selected gene expression profiles (n=59) for 6 non-tumor adrenal medulla (in white) and 60 pheochromocytomas (in black) tissues from Lopez-Jimenez et al. (2010). The color scale illustrates the relative level of gene expression: blue, highly expressed gene; red, low expressed gene. (B) A significant positive correlation was evidenced between the fold changes (FC) in expression of genes and corresponding cytosolic or membrane proteins of the exocytosis pathway. Linear regression (solid line) and 95% confidence bands (the regions delineated by dotted lines) are shown.

**Figure 7:**
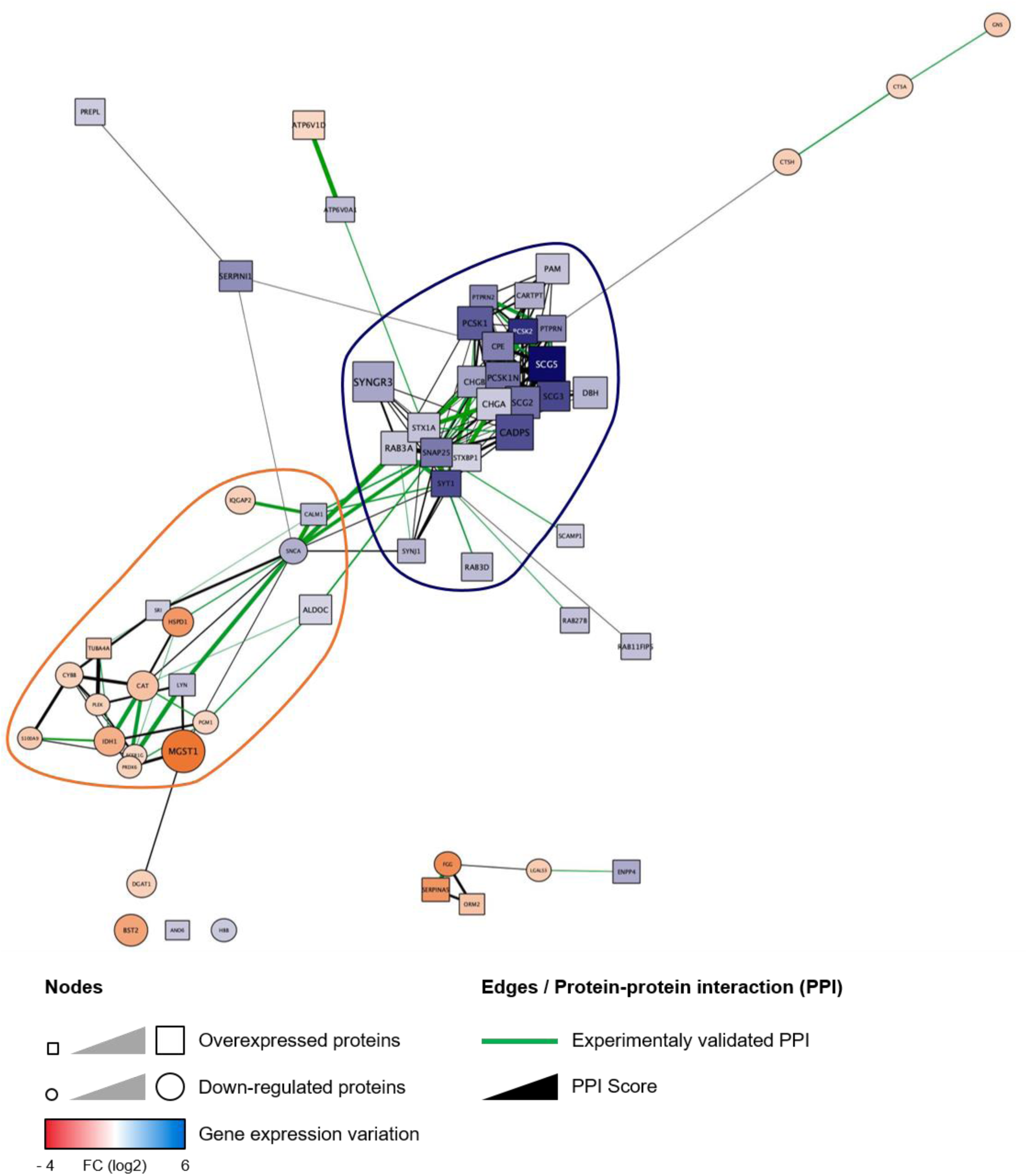
Protein-protein interaction (PPI) network analysis of the exocytic proteins modulated in Pheo at the transcriptomic and protein level. The PPI network was generated using Cytoscape. Overexpressed and downregulated proteins are represented by square and round nodes, respectively and the level of protein expression is proportional to their size whereas the mRNA expression variation is color coded as indicated. Therefore blue squares and red circles represent candidate for which both protein and mRNA are overexpressed or downregulated, respectively. The 2 first clusters identified by the CytoCluster plugin are outlined in blue (p=2.74×10^−9^) and red (p=3.65×10^−6^). Note that the variation observed at the protein and mRNA level goes in the same direction for 53 proteins out of 59 (90%).

Finally, to further validate our dataset, we analyzed the global expression change of some key overexpressed proteins present in the main exocytic cluster identified with Cytoscape. To do so, we performed a multiplexed MRM MS assay in total protein homogenates prepared from another independent cohort of 25 pairs of human Pheo and their matched non-tumor tissue (Croise *et al*, 2016). As MRM is a targeted MS approach that uses synthetic peptide reference standards, it is used to confirm and quantify the presence of proteins of interest on smaller amounts of sample with high sensitivity, which eliminates the need for fractionation (Keshishian *et al*, 2007). As observed for the subcellular fractions, using MRM MS we found that the expression of every protein identified in the PPI analysis was also significantly increased in Pheos compared with the matched adjacent non-tumor adrenal tissue (Figure 8).

**Figure 8:**
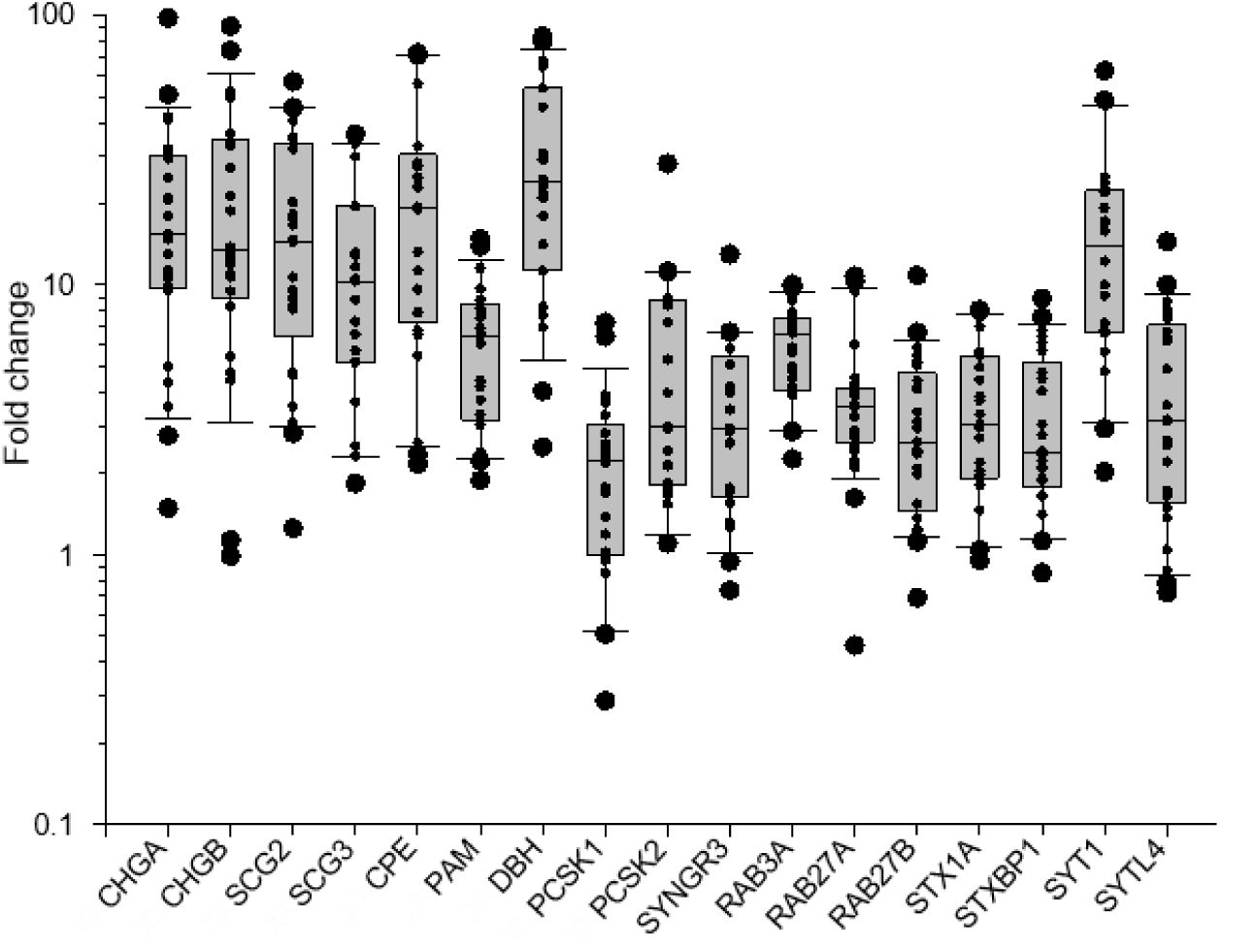
Global variation of key exocytic proteins expression in human Pheo tissue. Expression variation of the indicated proteins was quantified at the protein level by MRM-MS in 25 pairs of human Pheo normalized to their matched adjacent non-tumor tissue. Box- and-whisker diagrams illustrate the distribution of protein expression fold change (FC) for the 25 Pheos.

## Discussion

Dysfunction of hormones and neuropeptides secretion in NETs is a serious health issue. Patients with midgut primary carcinoids have increased serotonin and metabolite secretion, corresponding to higher metastatic tumor burdens (Onaitis *et al*, 2000). Hypersecretion of serotonin by carcinoid tumor from the gastro-intestinal tract can trigger carcinoid syndrome, which is associated with severe consequences such as flushing, diarrhea, bronchoconstriction and cardiac valvular disease (Onaitis *et al*., 2000). Acromegaly often result from excessive secretion of growth hormone by pituitary adenoma (Dineen *et al*, 2017). Excessive level of circulating catecholamines in patients with Pheo can trigger life-threatening medical problems such as cardiopathy and stroke (Y-Hassan & Falhammar, 2020; Zhang *et al*., 2017). Moreover, enhanced secretory activity of NET cells may develop over time with negative impact on prognosis. For example, silent pituitary adenoma can evolve into an active secreting adenoma whereas non-functional pancreatic tumors can become hormonally active, hence turning to a more aggressive tumor phenotype (Brown *et al*, 2006; Daems *et al*, 2009; Juhlin *et al*, 2019). Small cell lung cancer (SCLC) is a high-grade malignant cancer due to the progressive neuroendocrine nature of SCLC cells that secrete a variety of neuropeptides, together with growth factors that all dramatically accelerate the invasive growth by their autocrine action (Cuttitta *et al*, 1985; Song *et al*, 2003). Altogether, these few examples taken from a longer list clearly reveals that dysfunction of the secretory pathways in NETs can lead to severe clinical complications and can also impact the tumor development and prognosis. Today, a clear unmet need is to identify the cellular and molecular mechanisms triggering hypersecretory activity in NETs.

The aim of the present study was thus to uncover part of the mechanisms triggering catecholamine hypersecretion in human Pheo cells. To do so, we have used carbon fiber amperometry recordings to analyze catecholamine secretion on individual tumor cells cultured directly from freshly resected human Pheos. In parallel, we have analyzed the expression level of various proteins involved in calcium-regulated exocytosis by differential mass spectrometry methods applied on human Pheo tissues.

At the cellular level, several hypotheses which may not be mutually exclusive could explain the hypersecretory phenotype of Pheo: i) a leakage of catecholamines through the constitutive secretory pathway, ii) a global increase in the quantity of catecholamines in the secretory granules, iii) a simple mass effect, as the number of cells increases within the tumor and iv) a dysfunction of the calcium-regulated secretory pathway. Interestingly, our amperometric analysis has allowed us to identify the later possibility as a likely cause of this phenomenon. First, leakage of catecholamines in resting tumor cells does probably not occur as we never observed any spontaneous amperometric spike in the absence of cell stimulation (data not shown). Second, an increase of catecholamines in the secretory granules is rather unlikely since in most cases, quantal size of individual secretory events was not increased. Third, we showed that the number of exocytic events is significantly increased in single tumor chromaffin cells compared to non-tumor cells indicating that the cell proliferation within the tumor cannot be responsible alone for the hypersecretion phenotype. Moreover, by analyzing the spike kinetics, we have also observed that, for many patients, exocytic events tend to be faster. Altogether, our data clearly demonstrated that the regulation of calcium-regulated exocytosis is highly perturbed in tumor cells. To our knowledge, this is the first report analysing the catecholamine secretion using carbon fiber amperometry recording directly performed on human tumor cells from freshly resected pheochromocytomas.

The specific step of the exocytic pathway and which proteins might be involved in this amplified secretory activity remain to be explored in details. However, a significant increase in spike frequency can be consistent with an increase in efficiency of the different steps upstream of the fusion process including recruitment, docking and priming or with a greater calcium sensitivity of the exocytosis machinery. Faster kinetics of the exocytotic events might also reflect a direct effect on the fusion process. Through the differential mass spectrometry analysis, we have identified several proteins involved in the regulation of these various step of exocytosis, that are significantly over-expressed in the tumor tissue compared to the non-tumor tissue. Among them, some are known to affect the amperometric spike frequency and/or the kinetics of the spikes when their expression changes in chromaffin cells. This is the case for example of SNARE proteins (SNAP25, Syntaxin1), as well as for proteins regulating the SNARE complex (Synaptotagmin-1 and −7) or for the actin cytoskeleton organization (Rab27A or Annexin-A2) (Desnos *et al*, 2003; Fang *et al*, 2008; Gabel *et al*, 2015; Tawfik *et al*, 2021). Of note, among the core exocytotic machinery the calcium sensor synaptotagmin-1 was found to be among the most overexpressed in tumor cells (Figure 8), which could lead to an increase in calcium sensitivity and probability of release. Finally, it must be mentioned that at this stage, we cannot rule out an increase of the number of secretory granules in tumor cells as various specific soluble cargos of secretory granules, like different chromogranins or enzymes involved in hormone processing, were also found to be overexpressed. These observations are in line with previous reports indicating that chromogranins or chromogranin-derived peptides are highly expressed in Pheo (Guerin *et al*, 2010), and often found at higher levels in human fluids, which makes some of the proteins identified in our screen biomarker candidates.

To conclude, we have reported here that calcium-regulated exocytosis is deregulated in human Pheo cells and we have described tumor-associated expression changes of various key players of the exocytic pathway. The next challenge will be to understand how exactly these changes lead to the hypersecretion of catecholamines by the tumor. Moreover, a key unmet need is to find molecules able to prevent the catecholamine hypersecretion directly from the tumor cells. Of note, we have recently identified the somatostatin analog pasireotide (SOM230) as a *bona fide* inhibitor of Pheo hypersecretion (Streit *et al*, 2021), but the probable mechanism of action of this drug involves an inhibition of the cholinergic stimulation, which may prove to affect the activity of a large variety of secretory cells possessing nicotinic receptors. There is therefore a need to correct hypersecretion by identifying specific alterations in the secretory machinery whose potential candidates are identified in this study.

## Material and methods

### Subjects and samples

The medical files of patients with Pheo in 2 French centers (CHRU, Nancy and NHC, Strasbourg) between 2013 and 2020 were retrospectively reviewed. We collected initial diagnosis, including a clinical examination looking for hormonal-related symptoms and biological analysis. As recommended by the Endocrine Society clinical practice guideline published in 2014 (Lenders *et al*, 2014), Pheo genetic testing was proposed to identify germline mutations in the major susceptibility genes (SDHB, SDHC, SDHD, VHL, NF1, RET, TMEM127, MAX) using Sanger sequencing and multiplex ligation-dependent probe amplification (MLPA). Then, as recommended in the consensus statement published in 2017 (Group *et al*, 2017), next-generation-sequencing (NGS)-based diagnostic was carried out for more recent patients. Biological analysis comprised the measurement of metanephrine (MN) and normetanephrine (NMN) levels (in urine and/or plasma). When available, chromogranin A (CGA) measurements were also registered. Levels of free MN, NMN and CGA in plasma, as well as urinary levels of MN and NMN are presented as ratios normalized by the upper limits of normal (ULN). Plasma and urinary MN or NMN levels reaching two-fold the upper limit of the normal range and/or CGA exceeding the upper limit of the normal range was defined as the threshold of abnormal hormonal secretion (Lenders *et al*., 2020). The ULN of free MN and NM were 4.05 and 9.8 nmol/L in plasma and 1625 and 2620 nmol/24 h in urine, respectively. The upper reference limit for CGA was 100 mg/L. Catecholamine-producing phenotype of Pheo were categorized as previously described (Eisenhofer *et al*, 2005): adrenergic (AD) phenotype, when MN content exceeded 10% of the combined MN and NMN contents, or noradrenergic (NAD) phenotype when MN content remained below 10% of the combined MN and NMN contents. Pathological evaluation was reviewed, including tumor size, Ki-67 result and the PASS (Pheochromocytoma of the Adrenal Gland Scaled Score) as previously described (Thompson, 2002). Biological and clinical characteristics are summarized in Tables 1 and 2 whereas details per patient are described in Supplementary Table 1.

### Primary culture of human pheochromocytoma cells

Human tumor cells were cultured from freshly resected Pheo following surgery (Moog *et al*, 2018). In the operating room and immediately after the resection, the adrenal gland was cut longitudinally in two parts. Roughly a 1 cm^3^ piece of tumor tissue was dissected and immediately plunged into ice cold transport medium (Ca^2+^- and Mg^2+^-free Hank’s Balanced Salt Solution (CMF HBSS, Sigma) supplemented with 0.2% Fetal Bovine Serum (FBS, Gibco) and 1% penicillin/streptomycin (Sigma) or MACS Medium Tissue Storage solution (Miltenyi Biotec). Up to 3 hours after resection, the tumor sample was minced into 1 mm^3^ pieces in a dish containing CMF HBSS. Chunks were collected, centrifuged at 250 g for 5 min at room temperature and the pellet resuspended in 15 mL of complete medium (RPMI 1640 GlutaMAX™ (Gibco), 15% FBS, 1% penicillin/streptomycin). Red blood cells, debris and fat were separated from minced tissue by sedimentation for 15 min at room temperature. The supernatant was removed and 15 mL of complete medium were added to the pellet before centrifugation at 250 g for 5 min. Tumor pieces were resuspended in HBSS (in 10 times the tissue volume), containing 1.5 mg/mL of collagenase B (Roche) and 1 mg/mL of the protease dispase II (Gibco) and gently rocked for 45 min at 37°C. 5 min before the end of protease digestion, 0.1 mg/mL DNase I (Roche) was added to remove potential DNA clumps. Samples were left for a few minutes to sediment at room temperature and supernatant recovered (fraction 1). The pellet was resuspended in 5 mL of CMF HBSS and triturated for a couple of minutes to dislodge tumor cells from chunks. The remaining pieces were left few minutes to sediment and the supernatant recovered (fraction 2). Both fractions were centrifuged at 800 g for 5 min at room temperature. Cell pellets were resuspended in 2 mL of CMF HBSS. 4 mL of Red Blood Cell Lysis Buffer (Roche) were added before being gently rocked for 10 min at room temperature. The fractions were centrifuged at 500 g for 5 min and resuspended into complete medium. Cell viability and density were estimated under a microscope and 300 μL of cell suspension were seeded into type I collagen (Corning)-coated 35 mm dishes (MatTek). Cells were left to adhere overnight at 37°C in an incubator with water-saturated and 5% CO_2_ atmosphere. 2 mL of complete RPMI were added the following day and cells were used within two days.

### Carbon fiber amperometry

Human tumor cells from freshly resected pheochromocytoma were washed with Locke’s solution (140 mM NaCl, 4.7 mM KCl, 2.5 mM CaCl_2_, 1.2 mM KH_2_PO_4_, 1.2 mM MgSO_4_, 11 mM glucose, 0.01 mM EDTA and 15 mM HEPES, pH 7.5) and processed for catecholamine release measurements by amperometry as previously described (Houy *et al*, 2015; Tanguy *et al*, 2020). A carbon fiber electrode of 5 μm diameter (ALA Scientific Instruments) was held at a potential of +650 mV compared with the reference electrode (Ag/AgCl) and approached close to one cell. Secretion of catecholamines was induced by a 10 s pressure ejection of a 100 µM nicotine (Sigma) solution from a micropipette (Femtotips®, Eppendorf) positioned 10 μm from the cell and recorded over 60 s. The amperometric recordings were performed with an AMU130 amplifier (Radiometer Analytical), calibrated at 5 kHz, and digitally low-pass filtered at 1 kHz. Analysis of the amperometric recordings was performed as previously described with a macro (laboratory of Dr. R. Borges; http://webpages.ull.es/users/rborges/) written for Igor software (WaveMetrics), allowing automatic spike detection and extraction of spike parameters (Segura *et al*, 2000). The spike parameters analysis was restricted to spikes with amplitudes higher than 5 pA, which were considered as exocytic events. All spikes identified by the program were visually inspected. Overlapping spikes and spikes with aberrant shapes were discarded for parameters analysis. Quantal size (spike charge, Q) of each individual spike was measured by calculating the spike area above the baseline. Spike area is defined as the time integral of each transient current, Imax as the height of each spike, half-width as the width of each spike at half its height (*T*_1/2_) and *T*_peak_ as the spike rise time (Figure 1A).

### Tissue fractionation

Frozen tumor and matched non-tumor adjacent tissues were cut into small pieces (∼10 mm^3^), and 3 mL of homogenization buffer (0.25 M sucrose, 10 mM Tris pH 7.4, 100 units/mL of DNase I, 5 mM MgCl_2_, Complete protease inhibitor EDTA-free cocktail) was added per tissue sample, homogenized twice for 10 s and one time for 20 s using a polytron set at speed 4.0. Homogenates were filtered through a 180 μm nylon and brought to 3 mL with the homogenization buffer, if necessary. Light membranes were obtained by isopycnic centrifugation using discontinuous sucrose gradients in which samples brought to 1.4 M sucrose were layered by 1.2 M and 0.8 M sucrose. After centrifugation at 155,000 g for 2 hours at 4°C, the light membrane fraction located at the 0.8 M to 1.2 M sucrose interface was collected, snap-frozen in liquid nitrogen and stored at −80°C. The light membrane fraction is enriched with plasma membrane, Golgi, endosomes and secretory pathway associated membranes. The cytosol fractions were obtained by centrifuging 200 µL of crude homogenates at 150,000 g for 1 hour at 4°C. The supernatant was collected, snap-frozen and stored at −80°C.

The amounts of protein were determined using the bicinchoninic acid (BCA) assay according to the manufacturer’s instructions (Pierce).

### Mass spectrometry analysis

30 μg of samples (homogenates, cytosol or light membranes) was incubated in a denaturing buffer at final concentration of 7 M urea, 175 mM NH_4_HCO_3_, 8.75% v/v acetonitrile and incubated for 30 min at room temperature. Samples were then diluted to 1 M urea with water and digested with trypsin (Promega) overnight at 37°C at a ratio of 1 μg of trypsin per 10 μg of protein for homogenate and cytosol samples while for light membranes, the ratio was set at 1 μg of trypsin per 25 μg of proteins. Samples were reduced with 10 mM tris(2-carboxyethyl)phosphine (final concentration), incubated for 30 min at room temperature and then acidified to 0.5 M HCl. The samples were desalted using C18 96-well plates (3M Empore). The C18 eluates from homogenate samples were evaporated and stored at 4^°^C prior to MS analysis. The C18 eluates from light membrane and cytosol samples were collected in injection plates for strong cation exchange (SCX), dried by vacuum evaporation and stored at −20°C. To fractionate peptides by SCX chromatography, samples were solubilized with reconstitution buffer (0.2% v/v formic acid, 10% v/v acetonitrile for light membrane samples; 20 mM K_2_HPO_4_, 25% v/v acetonitrile for cytosol samples) and loaded on an SCX column. Three fractions were collected following elution using a salt gradient. At the end of each SCX fractionation batch, the collected fractions were stored at −80°C. Once the SCX fractionation was completed, the fractions were freeze-dried and then desalted. The eluates were divided equally into two 96-well plates; one plate for LC-MS/MS analysis and the other plate as a back-up. All plates were vacuum evaporated and stored at −20°C until analysis by LC-MS/MS. Samples were resuspended in 92.5/7.5 water/ACN+0.2% formic acid and analyzed by LC-MS/MS on a nanoAcquity UPLC (Waters) coupled to a Q-Exactive mass spectrometer (Thermo). Survey (LC-MS) and tandem mass spectrometry scans (MS/MS) were acquired in the same run. The resolution for the MS and MS/MS scans were 70,000 and 17,500, respectively. Peptide separation was achieved using a Waters nanoAcquity Symmetry UPLC Trap column (180 µm x 20 mm, 5 µm particle size) and a Waters nanoAcquity UPLC BEH300 analytical column (150 µm x 100 mm, 1.7 µm particle size). The mobile phases were (A) 0.2% formic acid in water and (B) 0.2% formic acid in acetonitrile. For each sample approximately 3.6 µg was loaded onto the trap column for 3 min at a flow rate of 10 µL/min. Peptides were separated using a linear gradient (92.5% A to 84% A) for 26 min, followed by (84% A to 75% A) for 14 min and a wash at 60% B for 2 min. The flow rate was 1.8 µL/min. Protein identification was accomplished using data acquired by LC-MS/MS. The MS/MS spectra were matched to the corresponding peptide sequences found in the UniProt human protein database using Mascot (Matrix Science, version 2.2.06.) software.

### Multiplexed multiple reaction monitoring (MRM) assay

For each of the selected proteins, five MRM-suitable peptides were selected by CellCarta’s in-house MRM Peptide Selection software. If possible, peptides that were detected by mass spectrometry were prioritized. The selected peptides were synthesized by JPT Peptide Technologies (Germany). Synthesized peptides were resolubilized in 25%/75% water/DMSO (v/v), pooled and diluted with 0.2% formic acid in water to a concentration of 200 pmol/mL. This peptide mix was used to develop the MRM assay. The optimal 2 transitions (combination of peptide precursor and fragment ion mass-to-charge ratio that are monitored by the mass spectrometer) per peptide were determined using selected reaction monitoring (SRM)-triggered MS/MS on a QTRAP 5500 instrument (AB Sciex) coupled to a nanoAcquity UPLC (Waters). An SRM transition was predicted for each peptide. The detection of this transition triggered the acquisition of a full MS/MS spectrum of the target peptide. The two most intense fragment ions (b or y fragment ions only) in the MS/MS spectrum for each acquired peptide were recorded by in-house developed software. The mass spectrometer collision energy (CE) was optimized for each transition with 5 different CE values automatically generated by in-house developed software. A solution containing all synthesized peptides at a concentration of 200 pmoL/mL was analyzed with the created MRM method. The two best peptides per proteins were selected to be monitored by the MRM assay.

The processed samples were resolubilized with 11 µL of a reconstitution solution containing 5 internal standard (IS) peptides each at 100 ng/mL. Eight (8) µL of material (∼10 µg) was analyzed by LC/MRM-MS. Peptide separation was achieved using a BioBasic C18 column (Thermo) (320 µm x 150 mm, 5 µm particle size). The mobile phases were (A) 0.2% formic acid in water and (B) 0.2% formic acid in acetonitrile. Peptides were separated using a linear gradient (92.5% A to 60% A) for 21 min, followed by a wash at 60% B for 2 min. The flow rate was 10 µL/min. The transition peak areas were integrated using Elucidator software (Rosetta Biosciences) in combination with software developed at CellCarta for automated MRM peak integration.

### Data processing and statistical analysis

For the analysis of the carbon fiber amperometry, the data was first standardized (mean=0 and variance=1). Data then was normalized by quantile method using *preprocessCore* R package. PCA analysis was performed using FactoMineR and factoextra R packages. Spearman’s rank order correlation was performed using *cor* function from stats R package and plotted using corrplot and ggplot2 R packages. For comparison between paired Tumor (T) and Non-Tumor (NT) patients, paired Wilcoxon test was performed. For comparison between Tumor (T) patients, non-paired Wilcoxon test was performed.

For the differential expression analysis by mass spectrometry, the intensity values for all detected components were log (base e) transformed with values < 0 replaced by 0. Intensity data was normalized to account for small differences in protein concentration between samples. A subset of the samples was used to create a reference sample against which all samples were then normalized. The normalization factors were chosen so that the median of log ratios between each sample and the reference sample over all the peptides was adjusted to zero. Intensities below Limit of Detection (LOD=100000) after normalization, were then linearly mapped to the range of (LOD/2, LOD) to avoid spurious large fold changes. Intensities above LOD were not changed. A two-way ANOVA model was used for the peptide level analysis and is defined as follows: I_ijk_=M+C_i_+S_j_+ε_ijk_ where I is the peptide intensity, M is the overall average intensity, C the ‘clinical group’ factor (matched non tumor and tumor), S the ‘patient’ factor that takes into consideration the ‘pairing’ nature of the data, and ε random error. FDR (False Discovery Rate) and q-value were calculated, based on the p-values obtained from the ANOVA model, using the Storey Tibshirani method to make multiple testing adjustments. Tukey’s HSD (Honestly Significant Difference) method is used to perform post hoc contrast among different groups.

One protein may have several identified and quantified peptides. The following ANOVA model, which is an extension of the two-way ANOVA used above in the peptide level analysis, takes this into consideration by introducing a ‘peptide factor’ in the model: I_ijkl_=M+C_i_+S_j_+P_k_+ε_ijkl_ where I is the protein intensity, M an overall constant, C the ‘clinical group’, S the ‘patient’ factor, and P the peptide factor. The number of the levels for P is protein-dependent, equal to the number of identified and quantified peptides for the protein.

For Multiple reaction monitoring (MRM) analysis, differential intensity (DI) ratios were calculated in pair wise comparisons for each transition as the median of the ratio of the normalized intensities of each group. Paired Student’s t-test was applied for the expression analysis. Protein-level statistics were also computed by linearly combining the transitions of a given protein into a single variable and then applying a t-test.

All differential expression analysis and data visualization were done using R. PCA analysis was done using *prcomp* function from stats R package and plotted using ggplot2 R package. Volcano plot was performed using *EnhancedVolcano* function from R package.

### Transcriptomic analyses

Gene expression data were retrieved from the GSE19422 dataset (https://www.ncbi.nlm.nih.gov/geo/query/acc.cgi?acc=GSE19422) (Lopez-Jimenez *et al*., 2010) which includes 6 normal adrenal medulla and 61 Pheos. Consistently with the proteomic analysis, the 23 paraganglioma samples were not considered. The GSM483021 Pheo sample was excluded from our analysis because its transcriptomic profile, as revealed by clustering and UMAP analyses (data not shown), is similar to that of normal samples suggesting that it is misidentified. The expression level of each gene was calculated as the geometric mean of the significantly differentially expressed probes identified with the GEO2R web tool using a Ben-jamin-Hochberg FDR correction (q-values < 0.05). Genes of the regulated exocytosis (#BPGO:0045055) and secretory granule (#CCGO:0030141) GO terms were then selected for further analysis. Unsupervised hierarchical clustering was performed with Bioconductor (v.3.13) and R software (v.4.1.0) using Pearson correlation as distance calculation and complete linkage. The Fold change (FC) for each gene was calculated from the median expression values for the normal and Pheo groups and only those with a FC greater than or equal to 2 were retained for further analysis.

The correlation between the expression level of common genes and proteins was evaluated in both cytosolic and membrane fractions using Spearman correlation (GraphPad Prism v.8.1). In order to highlight functional clusters, an integrative analysis was conducted with Cytoscape software (v.3.8.2, (Shannon *et al*., 2003)) using expression data and protein interaction data retrieved from STRING-DB (v.11.5, https://string-db.org/, (Szklarczyk *et al*, 2021)). An edge-weighted force directed layout has been applied to the interaction network. Gene/protein clusters were identified using the CytoCluster plugin for Cytoscape using the default settings except for minimum size: 6, minimum density: 0.25, Edge weights: Feature Distance, Node penalty:3.

## Ethics approval and consent to participate

The present study used the data and the human biological material of the biological collection “Approche moléculaire des tumeurs corticosurrénaliennes” which was agreed by the “Comité de Protection des Personnes Est III” ethical advisory committee, and was conducted according to currently accepted ethical guidelines, including informed written consent approval signed by all patients prior to inclusion.

## Funding and acknowledgements

This work was financially supported by ITMO Cancer AVIESAN (Alliance Nationale pour les Sciences de la Vie et de la Santé, National Alliance for Life Sciences & Health) within the framework of the Cancer Plan to SG and LB (Single Cell 2018 N° 19CS004-00); by the University of Strasbourg Institute for Advanced Study (USIAS) for a Fellowship, within the French national programme “Investment for the future” (IdEx-Unistra) to SG; by the Conseil Régional de Normandie to CD; by grants from the Agence Nationale pour la Recherche (“SecretoNET”, N° ANR-16-CE17-0022-01) and from the Ligue contre le Cancer to SG (CCIR Grand-Est) and to CD (Comité Normand); by a fellowship from la Fondation pour la Recherche Médicale (FRM; FDM201806005916) to SM. INSERM is providing salary to SG and NV.

---

**Supplementary Figure 1:**
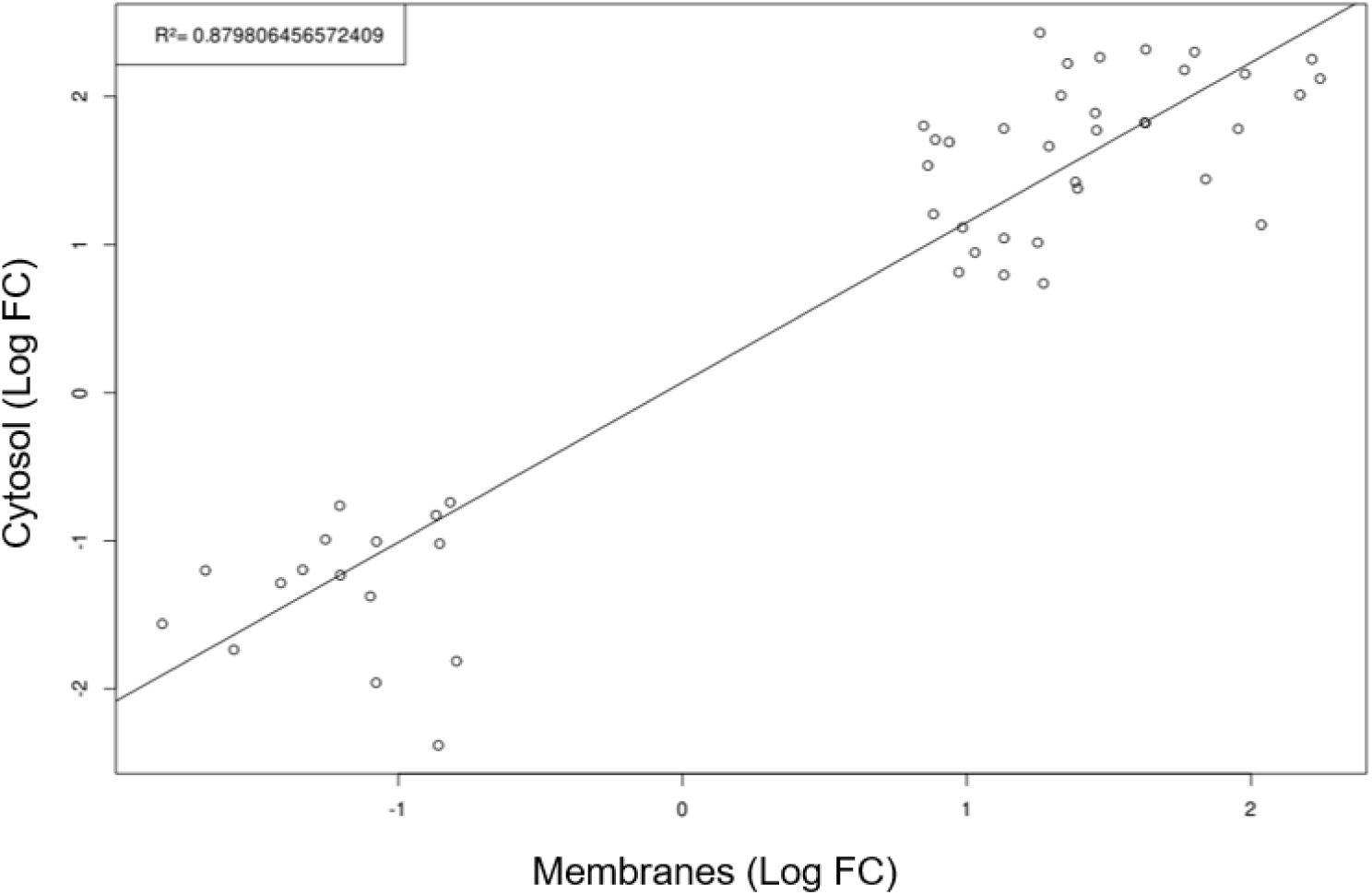
Proteins found in common in the membrane and the cytosolic fractions vary in the same direction. Among the 166 identified proteins whose expression significantly changes more than 2 fold, 50 are found in both cytosol- and membrane-enriched fractions. Linear regression analysis between the membrane-fold changes and the cytosolic-fold changes shows a positive correlation for both down-regulated and up-regulated proteins.

**Supplementary Figure 2:**
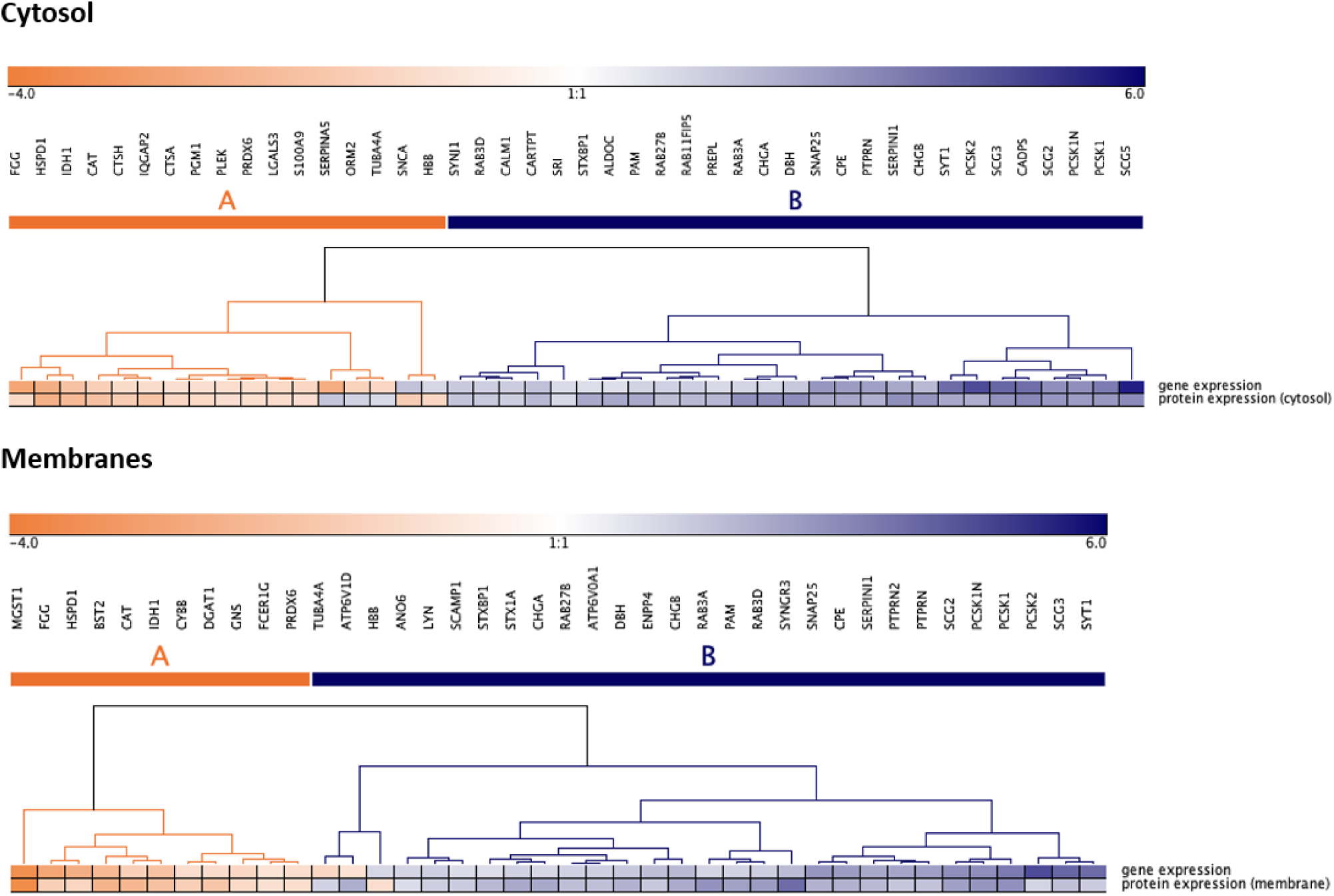
Comparison of genes and proteins of the exocytic pathway deregulated in Pheo. The 59 common differentially expressed genes (GSE19422) and proteins (this study) are selected from the exocytosis pathway. Unsupervised hierarchical clustering highlights two clusters of up- and down-regulated genes and proteins. Except for 5 (A) and 3 (B) genes/proteins, expression of all the genes (top row) and their proteins (bottom row) varies in the same direction. The color scale illustrates the over- (blue) or under- (red) expression of genes and proteins in Pheo compared to non-tumor tissue.

**Supplementary Table 1:**
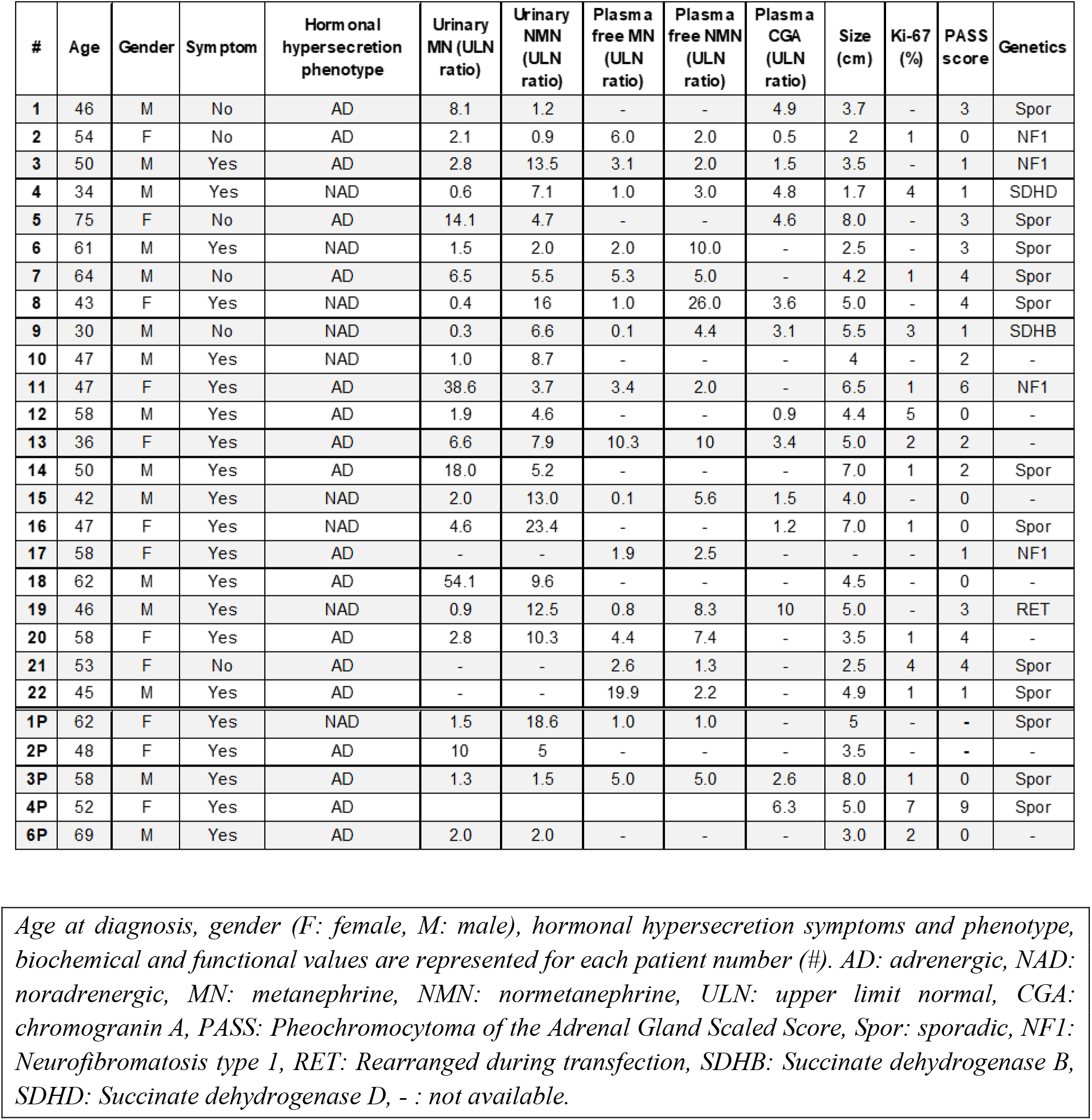
Clinical, biochemical, and functional characteristics of the 27 patients with Pheo evaluated in this study.

**Supplementary Table 2:**
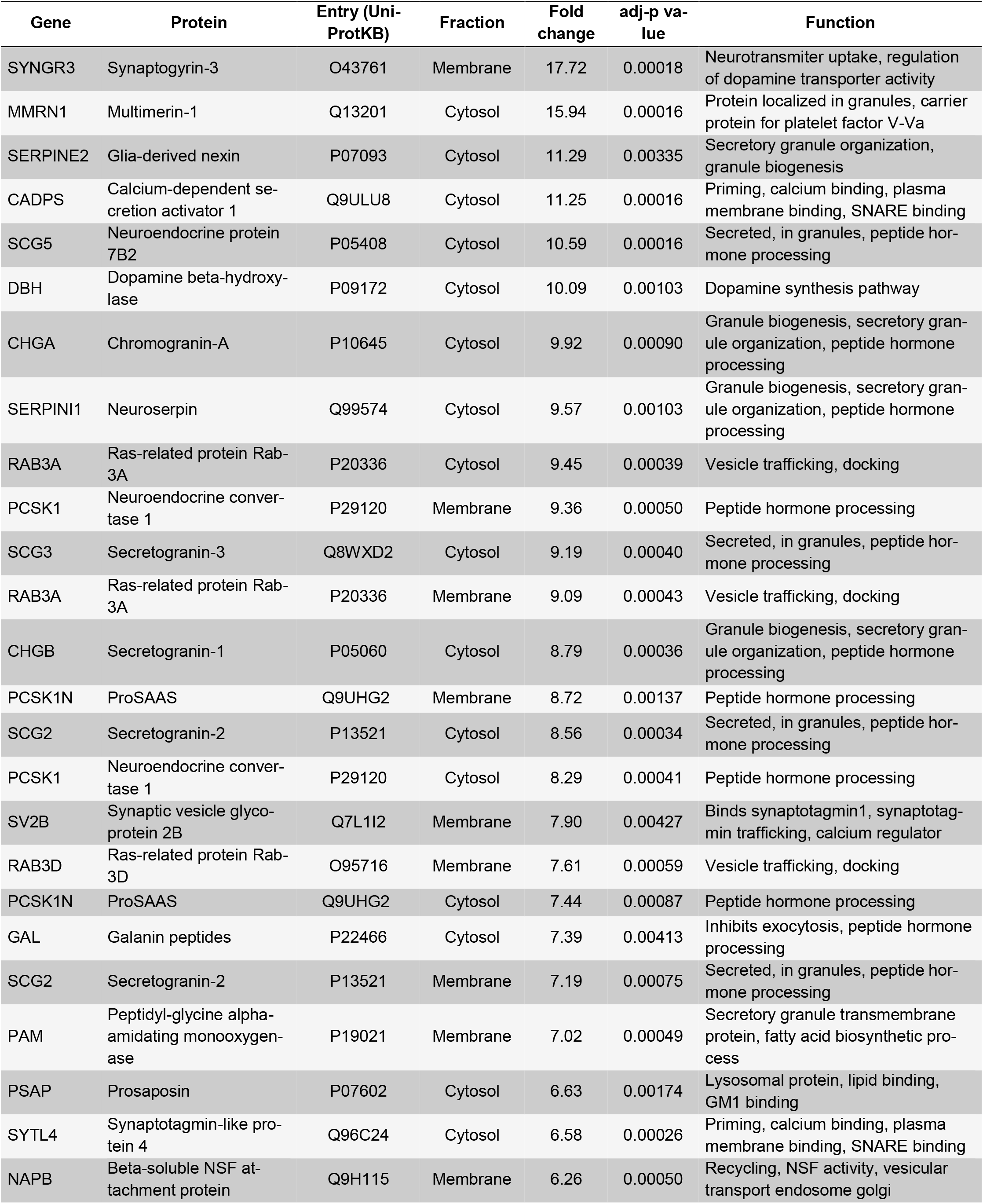
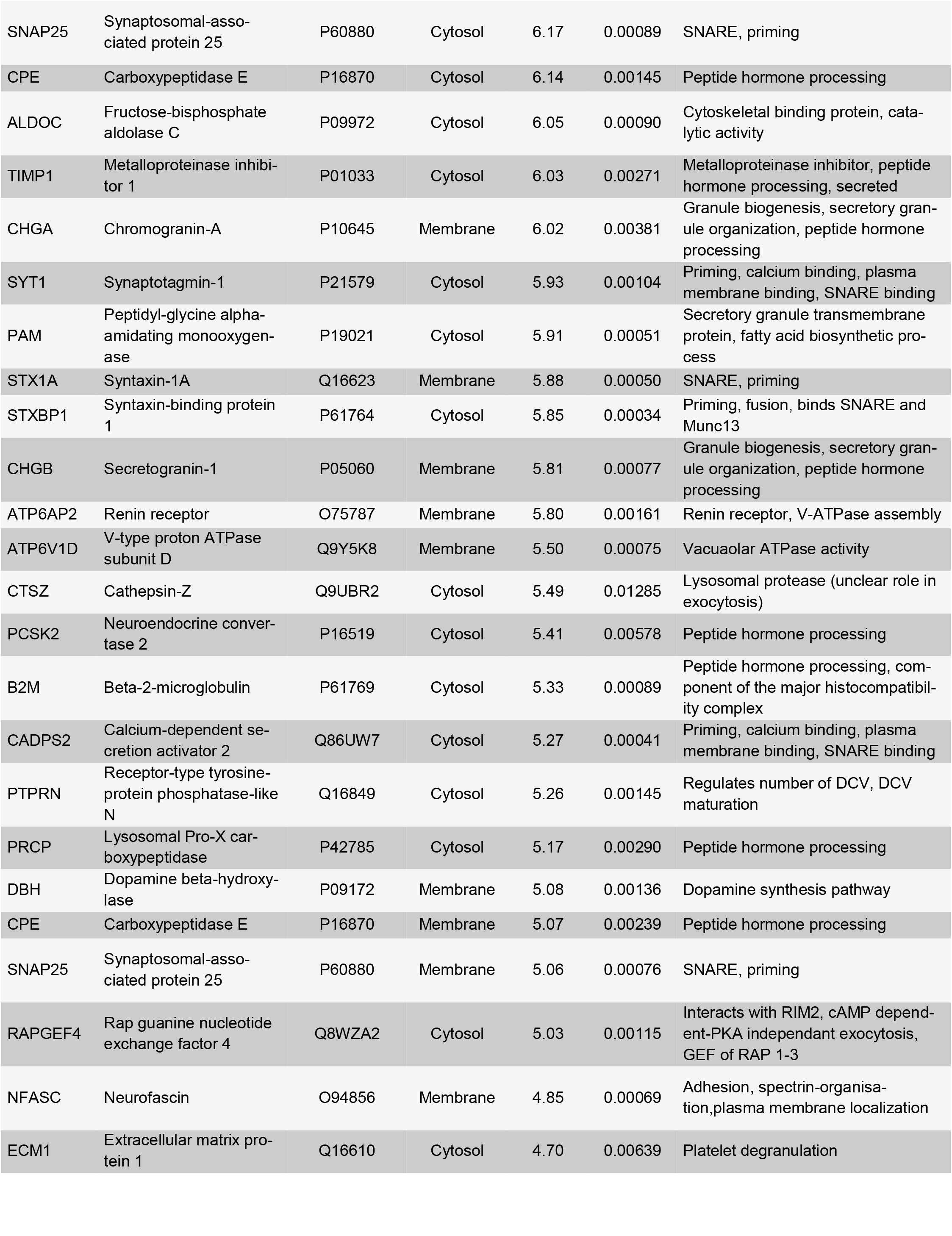
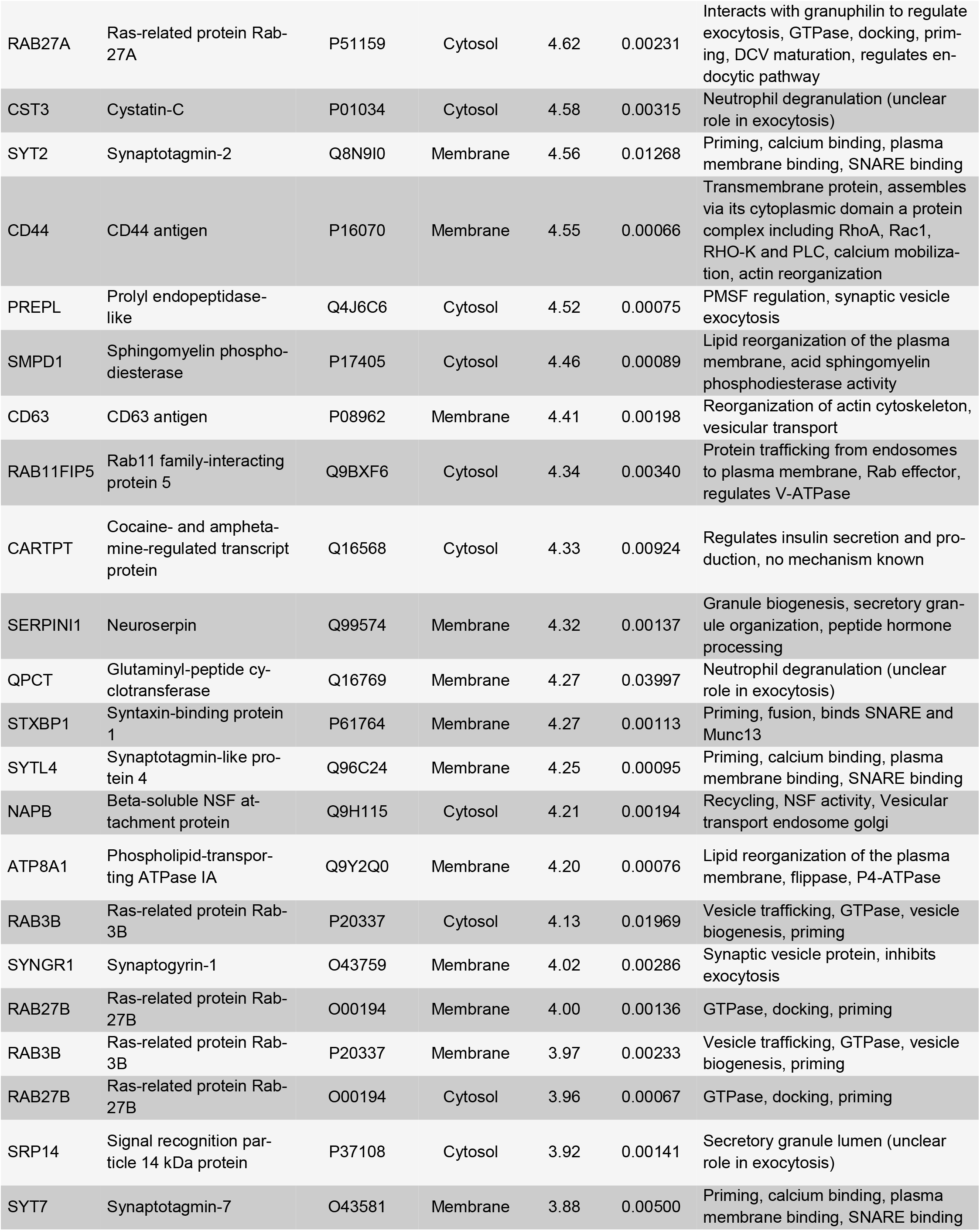
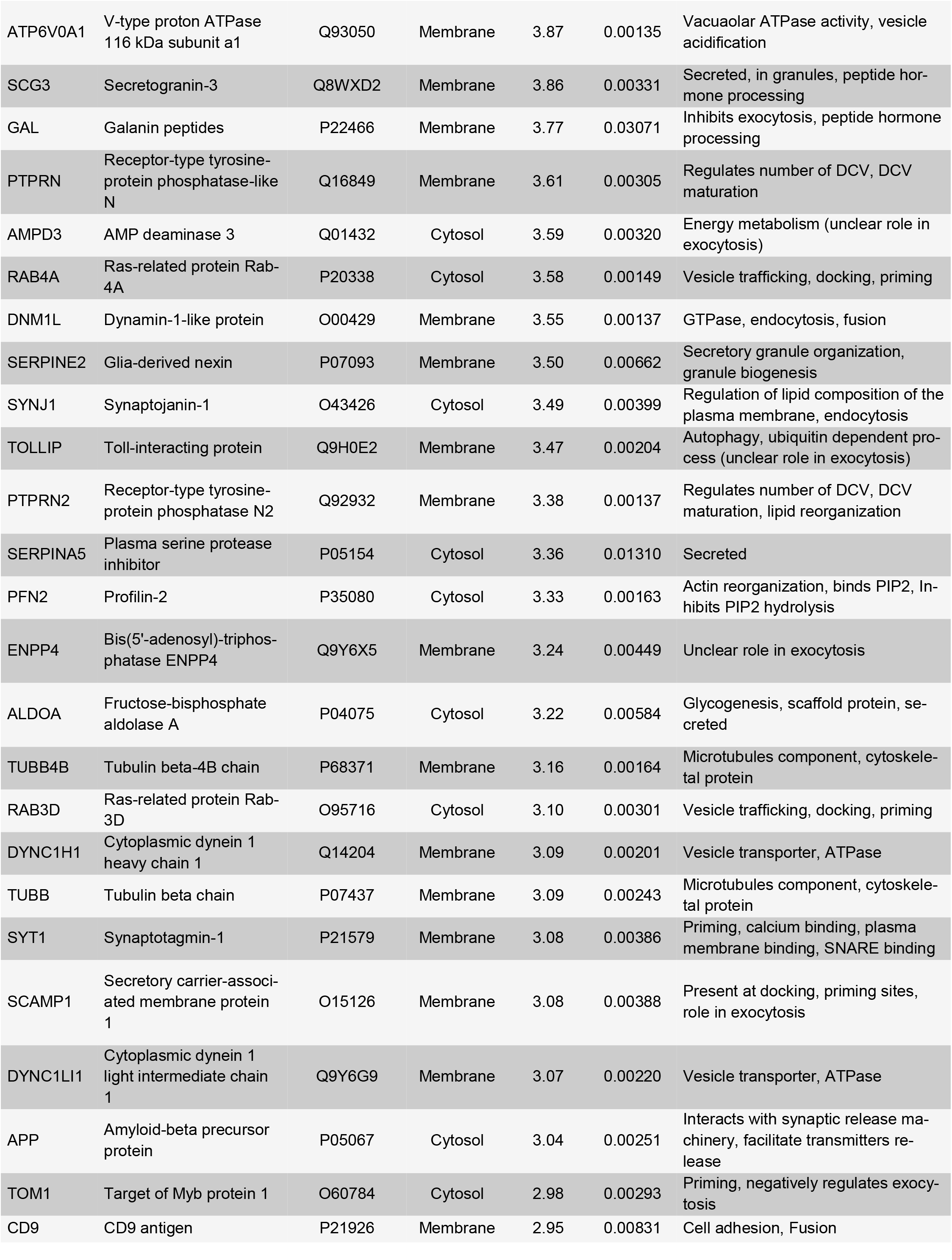
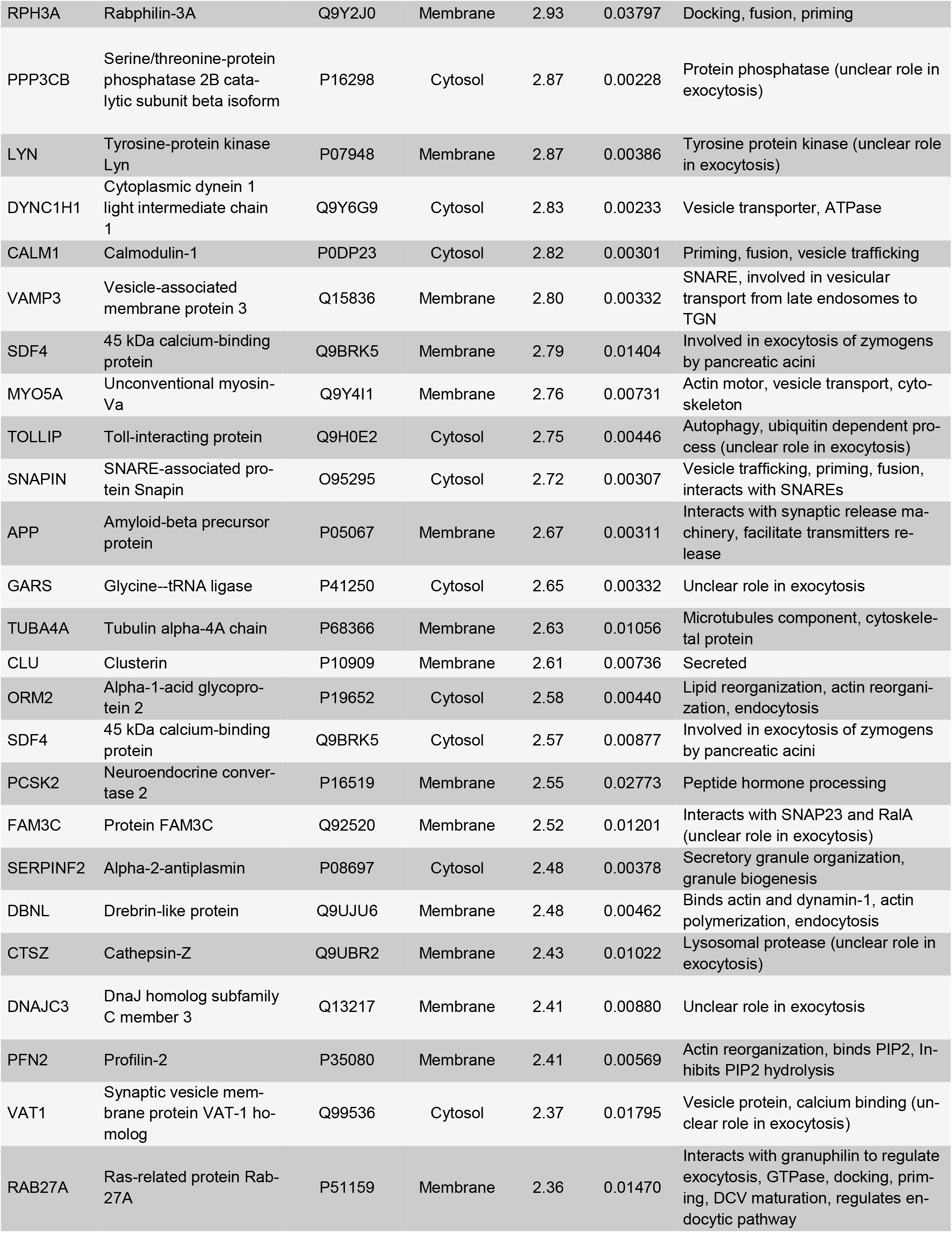
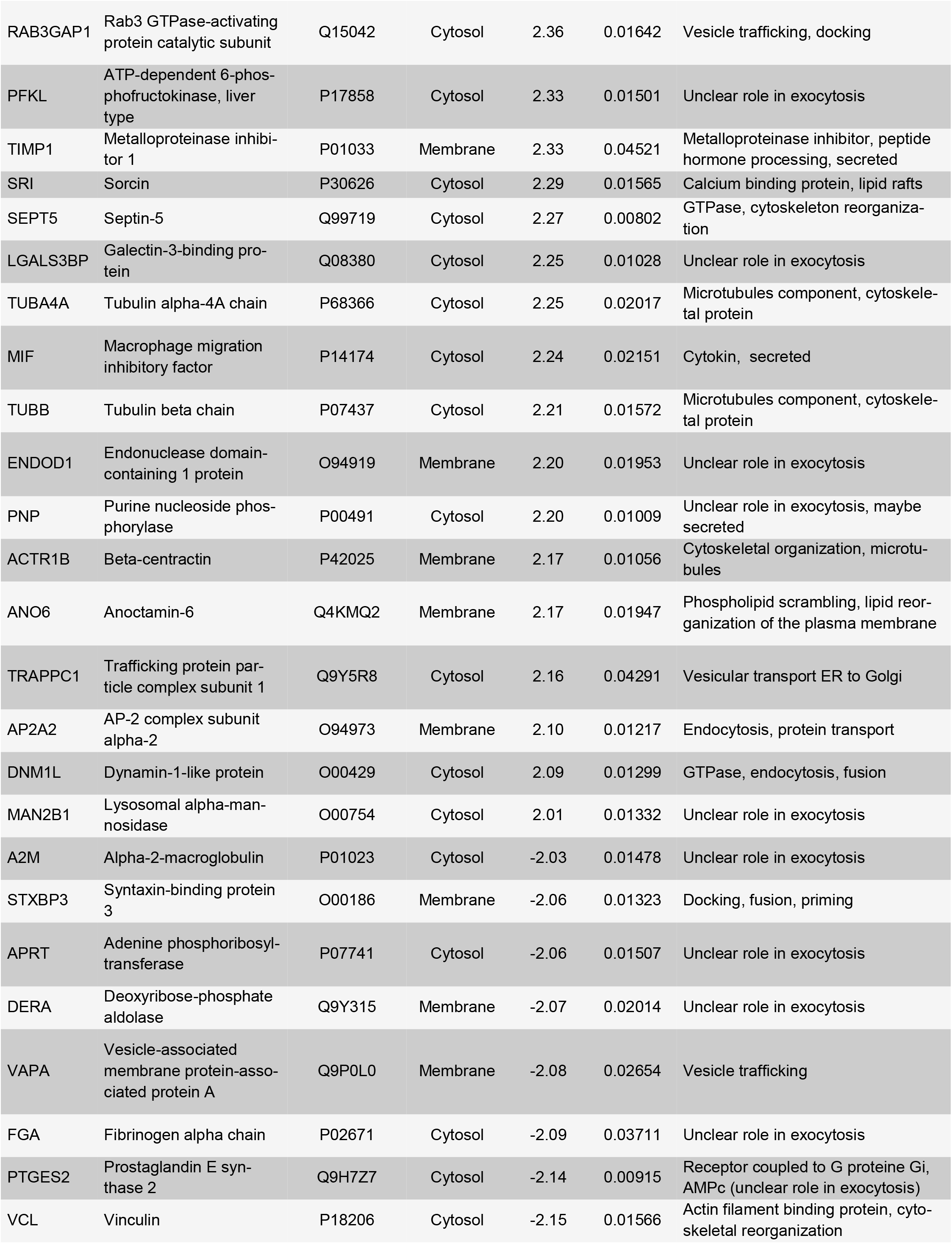
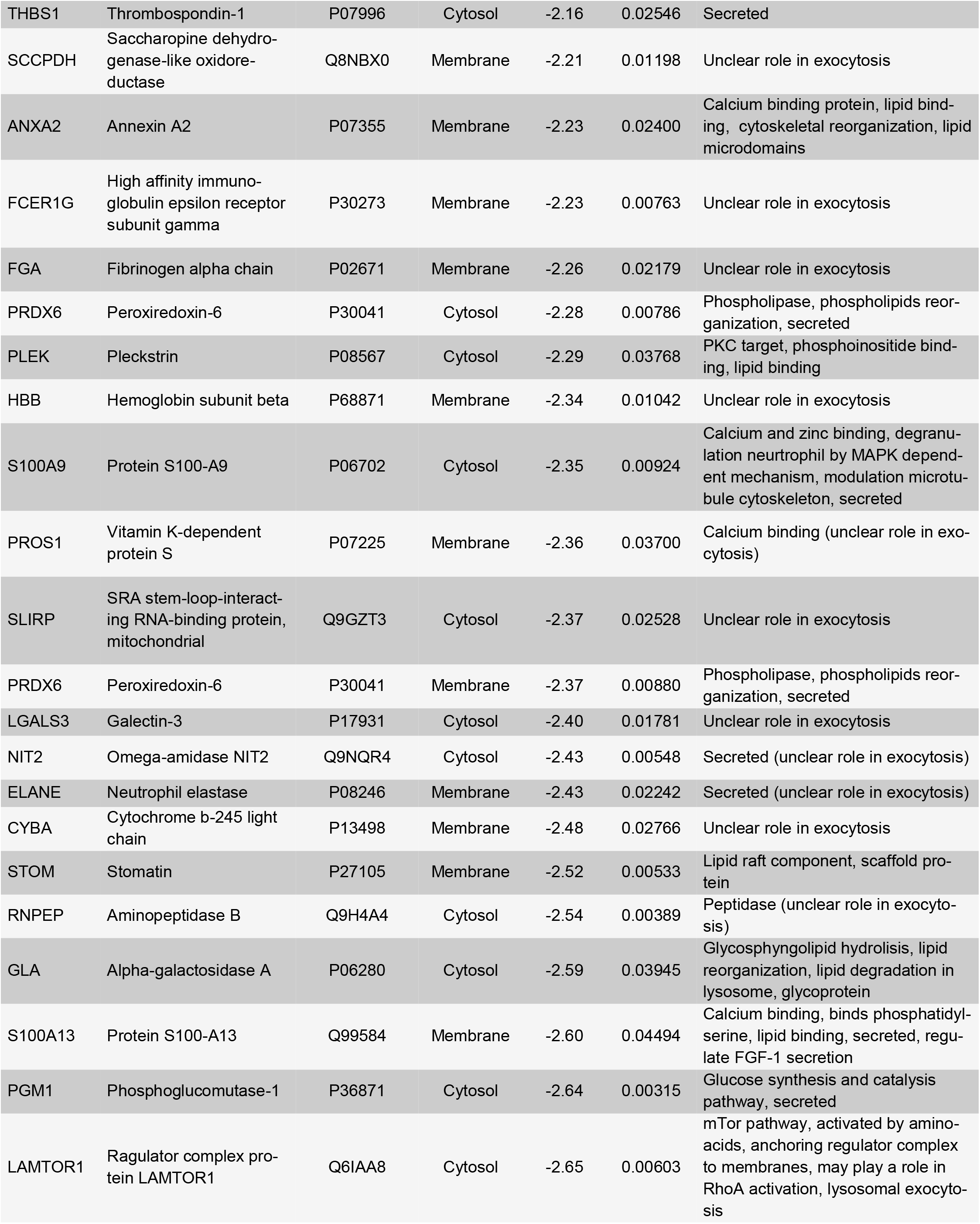
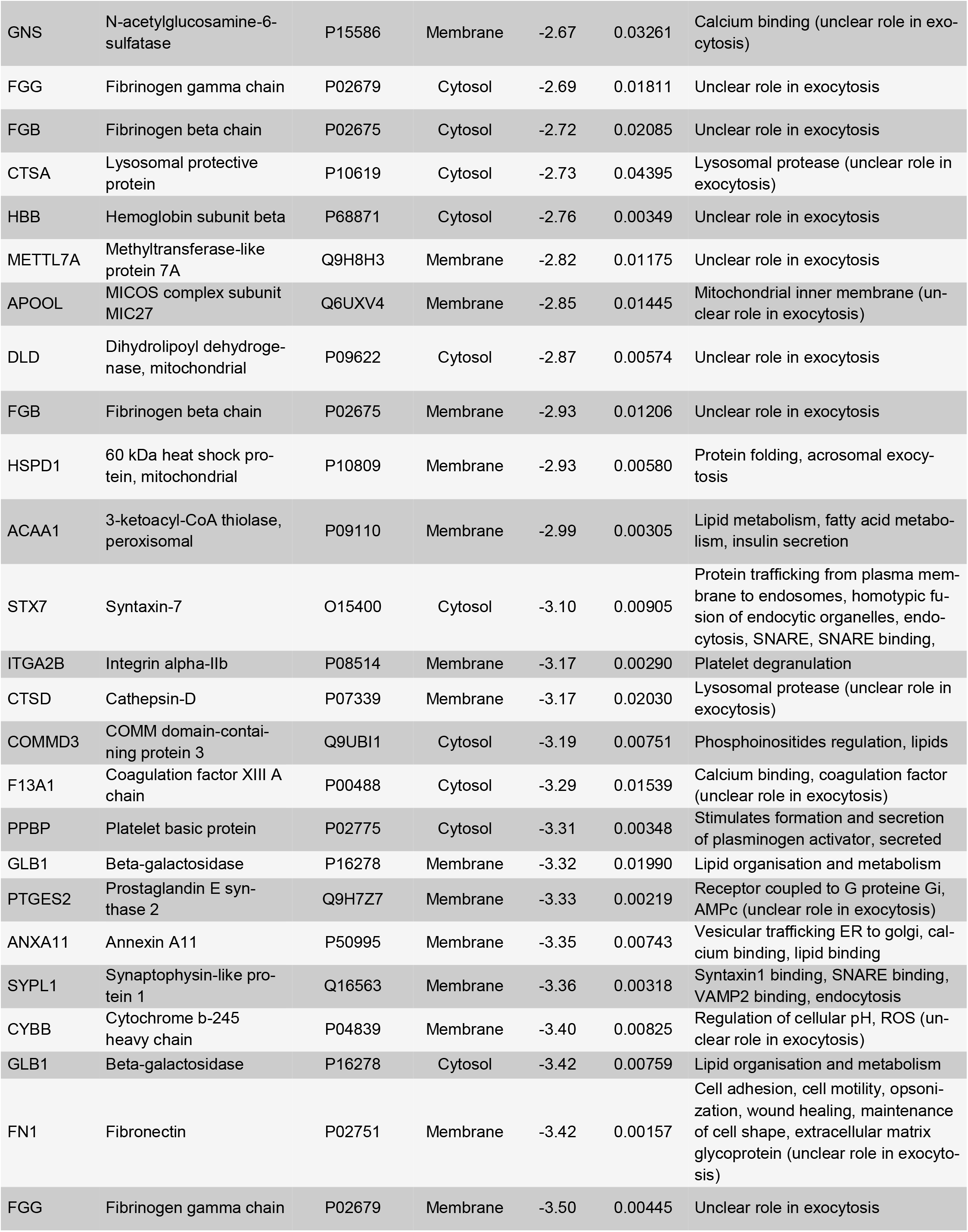
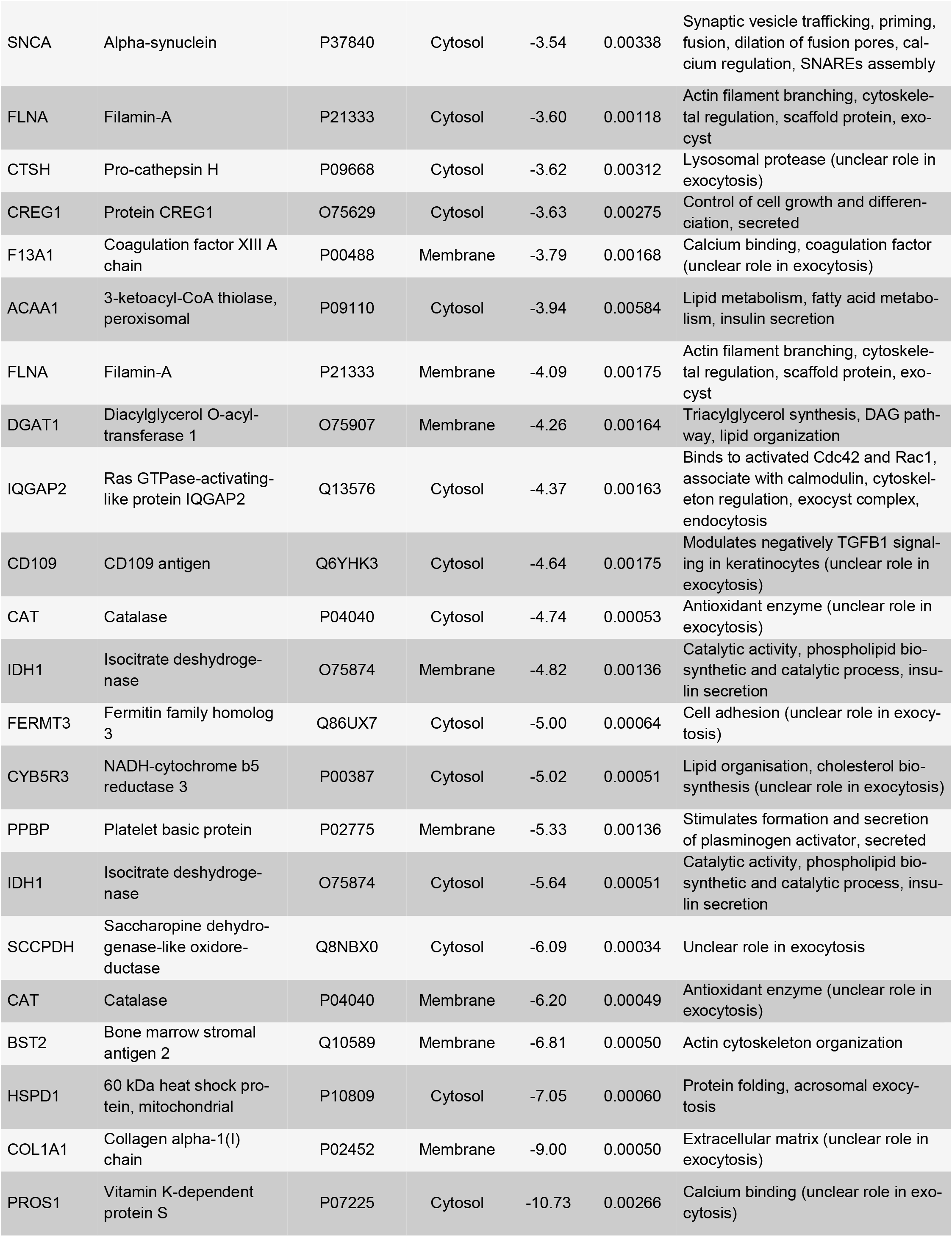
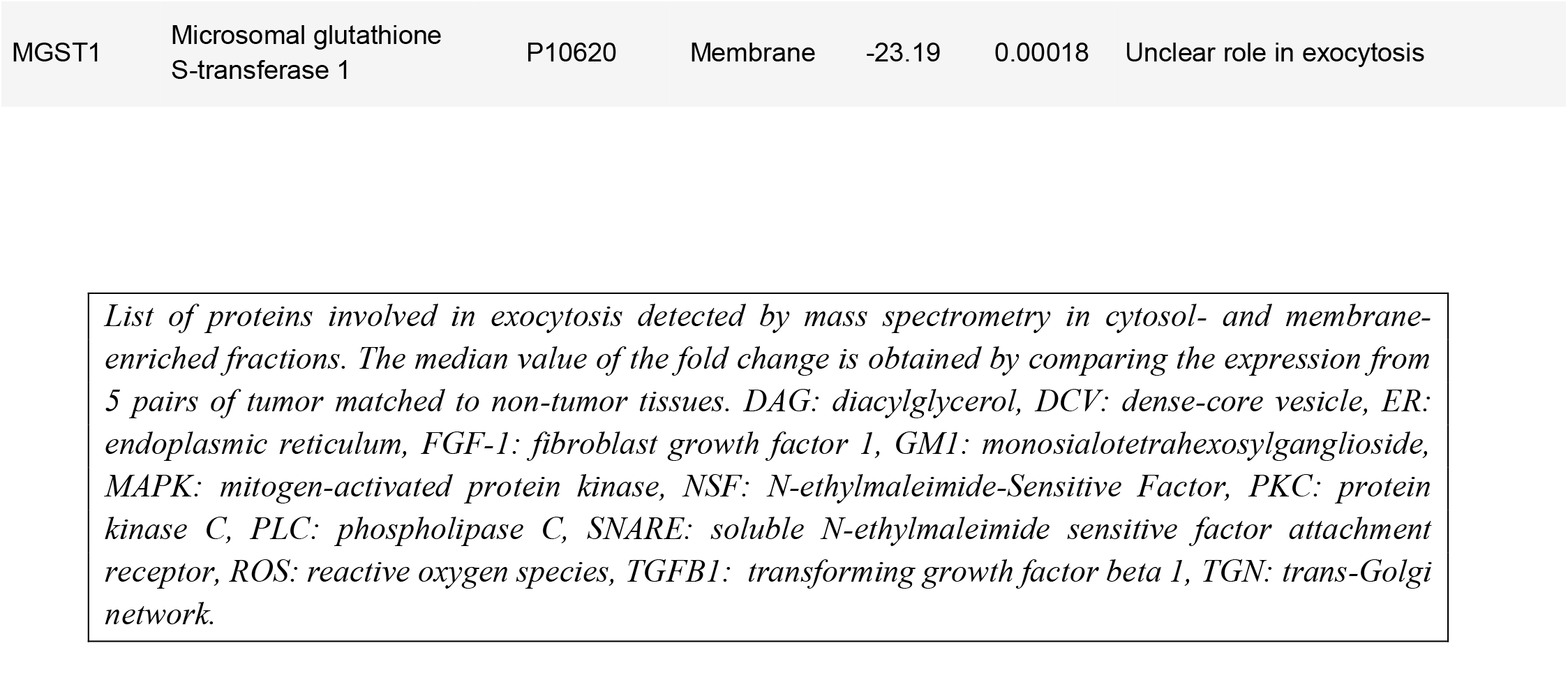
List of up- and down-regulated proteins in Pheo by comparison with the adjacent non-tumor tissue.

